# Identifying and correcting for misspecifications in GWAS summary statistics and polygenic scores

**DOI:** 10.1101/2021.03.29.437510

**Authors:** Florian Privé, Julyan Arbel, Hugues Aschard, Bjarni J. Vilhjálmsson

## Abstract

Publicly available genome-wide association studies (GWAS) summary statistics exhibit uneven quality, which can impact the validity of follow-up analyses. First, we present an overview of possible misspecifications that come with GWAS summary statistics. Then, in both simulations and real data analyses, we show that additional information such as imputation INFO scores, allele frequencies, and per-variant sample sizes in GWAS summary statistics can be used to detect possible issues and correct for misspecifications in the GWAS summary statistics. One important motivation for us is to improve the predictive performance of polygenic scores built from these summary statistics. Unfortunately, due to the lack of reporting standards for GWAS summary statistics, this additional information is not systematically reported. We also show that using well-matched LD references can improve model fit and translate into more accurate prediction. Finally, we discuss how to make polygenic score methods such as lassosum and LDpred2 more robust to these misspecifications to improve their predictive power.

## 1 Introduction

Contrary to individual-level genotypes and phenotypes, summary statistics resulting from genome-wide association studies (GWAS) are widely available, and very large sample sizes can be obtained through meta-analyses (Yengo *et al*., 2022). GWAS summary statistics have been extensively used to derive polygenic scores (PGS), perform fine-mapping, and estimate a range of key genetic architecture parameters (Pasaniuc and Price, 2017; Privé et al., 2021; Chen et al., 2021). However, GWAS summary statistics come with uneven imputation accuracy. There is also heterogeneity in the information made available in these summary statistics; for example, per-variant imputation INFO scores, sample sizes and allele frequencies are often missing. Moreover, many methods based on summary statistics use Bayesian models and iterative algorithms, which can be sensitive to model misspecifications (Walker, 2013; Miller and Dunson, 2019).

We present an overview of possible misspecifications that come with GWAS summary statistics in Table 1. First, the total sample size used can be misestimated, which would result in a biased estimation of the SNP heritability. For example, the total effective sample size is overestimated when computed from the total number of cases and controls from a meta-analysis of binary outcomes (Grotzinger *et al*., 2021); using BOLT-LMM summary statistics can result in an increased effective sample size (Loh *et al*., 2018; Gazal et al., 2018); and using SAIGE on binary traits with a large prevalence can result in a reduced effective sample size (Zhou *et al*., 2018; Mbatchou et al., 2021). Second, per-variant sample sizes can vary substantially and be much smaller than the total sample size when meta-analyzing GWAS summary statistics from multiple cohorts with different sets of variants (Wang *et al*., 2021). Using very different per-variant sample sizes can be problematic for models implicitly assuming that summary statistics have all been derived from the same individuals (Zhu and Stephens, 2017; Zhou and Zhao, 2021). Third, many genetic variants are imputed, with association statistics being reported for the allele dosages instead of the true alleles, which can lead to some bias in the summary statistics. Fourth, the imputation quality of each variant (e.g. INFO scores), often used in a quality control (QC) step in statistical genetics analyses, can be severely misestimated (often overestimated) when computed from multi-ancestry individuals instead of a more homogeneous subset. Fifth, errors such as allele inversions can be present in the summary statistics; QC is particularly important here. Sixth, many follow-up analyses require using linkage disequilibrium (LD) from a reference panel. There can be some mismatch between the GWAS summary statistics and the LD reference used, which can e.g. lead to suboptimal predictive performance for polygenic scores.

**Table 1:**
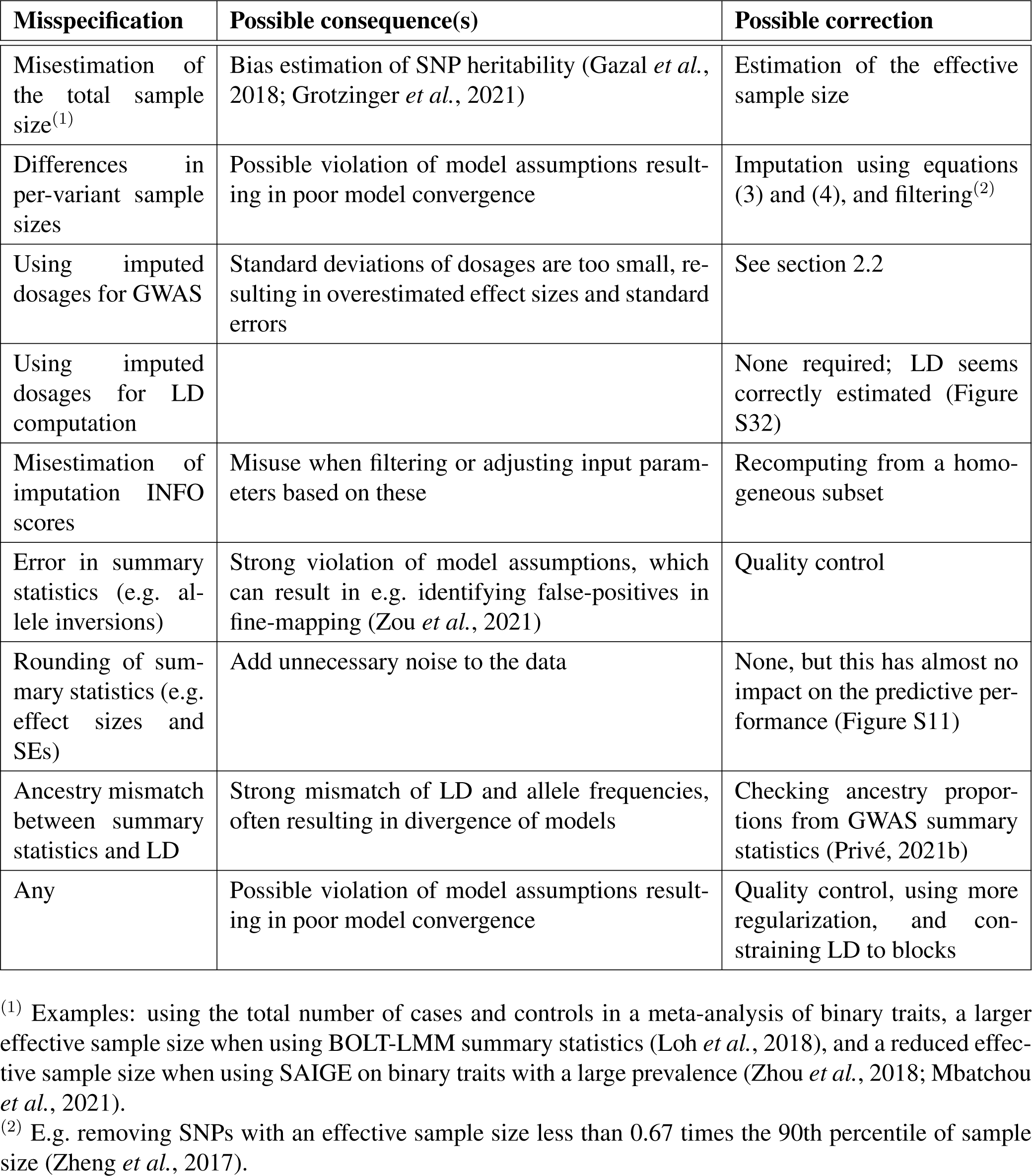
Overview of possible misspecifications when using GWAS summary statistics, along with possible consequences and corrections.

Here we investigate some of these misspecifications, and propose adjustments to improve the predictive performance of polygenic scores derived from GWAS summary statistics. We approach this from three different angles. First, based on additional summary information such as the imputation INFO scores and allele frequencies from the GWAS summary statistics, we refine our previously proposed QC (Privé *et al*., 2020b). This QC consists in comparing standard deviations (of genotypes) inferred from GWAS summary statistics with the ones computed from a reference panel. This is useful to check that the input parameters used are consistent with one another. This was particularly important for LDpred2-auto, which directly estimates two key model parameters, the SNP heritability and polygenicity, from the data (Privé *et al*., 2020b). Here we further show that standard deviations of imputed genotypes (allele dosages) are lower than the expected values under Hardy–Weinberg equilibrium. Second, we investigate possible adjustments to apply to the input parameters of PGS methods using summary statistics, namely the reference LD (linkage disequilibrium) matrix, the GWAS effect sizes, their standard errors and corresponding sample sizes. For example, we show that GWAS effect sizes computed from imputed dosages are larger in magnitude compared to if computed from true genotypes. Third, we introduce two new optional parameters in LDpred2-auto to make it more robust to these types of misspecification. We focus our investigations on LDpred2 and lassosum for two reasons. First, multiple studies have shown that LD-pred2 and lassosum rank among the best methods for single-trait polygenic prediction (Mak *et al*., 2017; Privé *et al*., 2020b; Pain et al., 2021; Kulm et al., 2021). Second, we reimplement and use a new version of lassosum, called lassosum2, that uses the exact same input parameters as LDpred2, which makes it easy for us to test the different QCs and adjustments presented here.

## 2 Results

### 2.1 Misspecification of per-variant GWAS sample sizes

We design GWAS simulations where variants have different sample sizes, which is often the case when meta-analyzing GWAS summary statistics from multiple cohorts with different sets of variants (Wang *et al*., 2021). Using 40,000 variants from chromosome 22 (Methods), we simulate quantitative phenotypes with a heritability of 20% and 2000 causal variants. We then divide the 40,000 variants into three groups: for half of the variants, we use 100% of 300,000 individuals for GWAS, but use only 80% for one quarter of the variants and 60% for the remaining quarter. We then run C+T, lassosum, lassosum2 (Methods section 4.4), LDpred2-inf, LDpred2(-grid), and LDpred2-auto (Privé *et al*., 2019; Mak et al., 2017; Privé et al., 2020b) by using either the true per-variant GWAS sample sizes, the total sample size for all variants, or per-variant sample sizes imputed using equation (3). Note that we initially included PRS-CS and SBayesR in our comparison (Ge *et al*., 2019; Lloyd-Jones et al., 2019). However, results for SBayesR always diverged (stopping with an error), and the overlap with the LD reference provided for PRS-CS was too small. When providing true per-variant GWAS sample sizes, squared correlations between the polygenic scores and the simulated phenotypes are of 0.123 for C+T, 0.161 for lassosum, 0.169 for lassosum2, 0.159 for LDpred2(-grid), 0.140 for LDpred2-auto, and 0.141 for LDpred2-inf (Figure 1, averaging over 10 simulations). Results when using imputed sample sizes are quite similar to when using the true ones. Note that C+T does not use this sample size information. When using the total GWAS sample size instead of the per-variant sample sizes, predictive performance slightly decreases to 0.157 for lassosum and to 0.163 for lassosum2, but dramatically decreases for LDpred2 with new values of 0.134 for LDpred2-grid, 0.119 for LDpred2-auto, and 0.123 for LDpred2-inf (Figure 1). This extreme simulation scenario shows that LDpred2 can be sensitive to the misspecification of per-variant GWAS sample sizes, whereas lassosum (and lassosum2) seems little affected by this.

**Figure 1:**
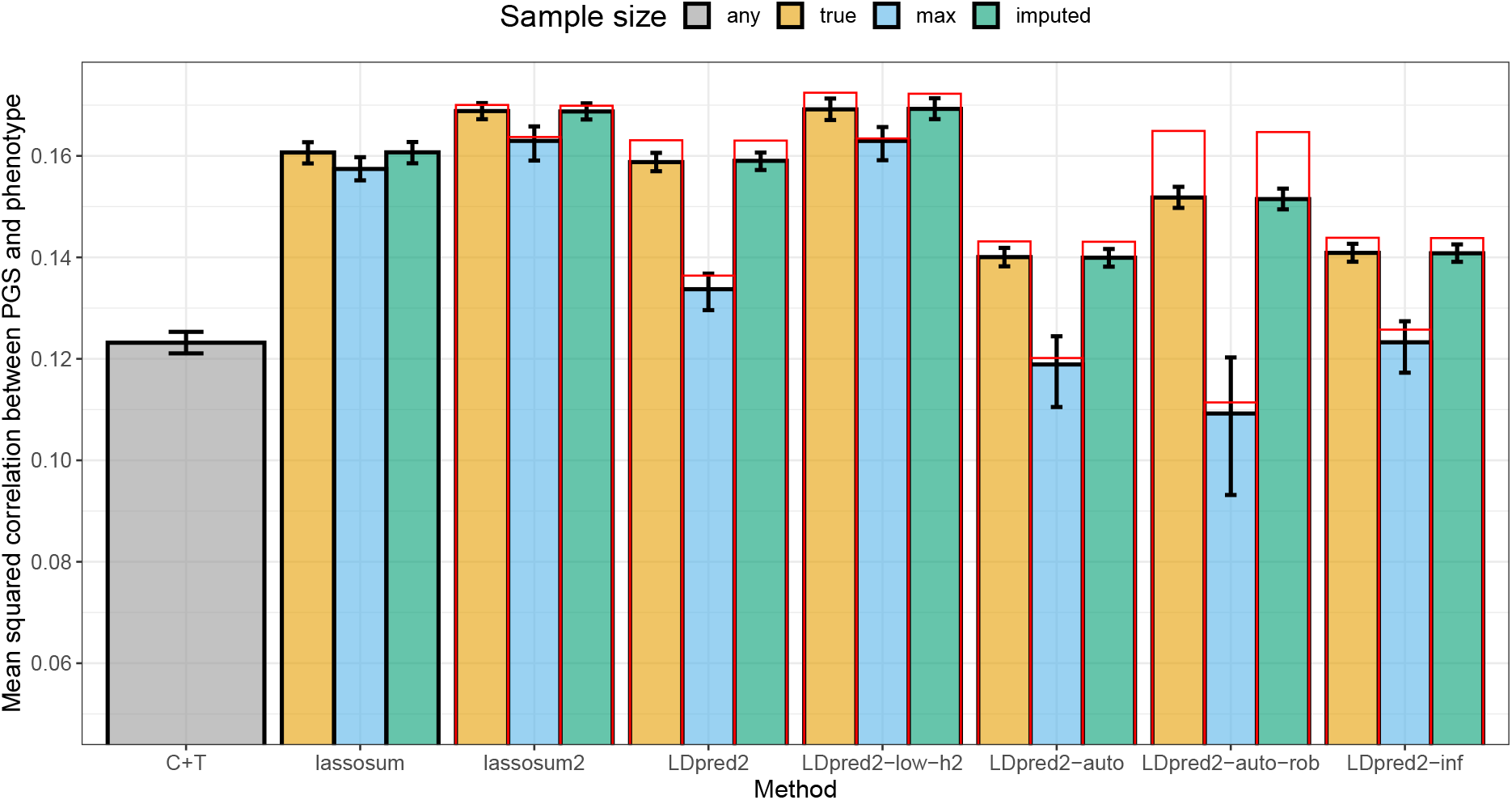
Results for the simulations with sample size misspecification, averaged over 10 simulations for each scenario. Reported 95% confidence intervals are computed from 10,000 non-parametric bootstrap replicates of the mean. The GWAS sample size is either “true” when providing the true per-variant sample size, “max” when providing instead the maximum sample size as a unique value to be used for all variants, “imputed” (cf. equation (3)), or “any” when the method does not use this information (the case for C+T). Red bars correspond to using the LD with independent blocks (Methods section 4.6); this strategy cannot be tested straightforwardly for C+T and lassosum.

We conduct further investigations to explain the results of figure 1. First, the results for LDpred2-auto are similar to LDpred2-inf because it always converges to an infinitesimal model (*p* = 1) in these simulations. To overcome this limitation, we introduce two new parameters to make LDpred2-auto more robust, and refer to this as “LDpred2-auto-rob” here (Methods section 4.5). The first parameter prevents *p* from diverging to 1, while the second shrinks the off-diagonal elements of the LD matrix (a form of regularization). Second, for lassosum2, results for a grid of parameters (over *λ* and *δ*) are quite smooth compared to LDpred2 (Figures S2 and S3). In these simulations with misspecified per-variant sample sizes, it seems highly beneficial to use a small value for the SNP heritability hyper-parameter *h*^2^ in LDpred2, e.g. a value of 0.02 or even 0.002 when the true value is 0.2 (Figure S3). Indeed, using a small value for this hyper-parameter induces a larger regularization (shrinkage) on the effect sizes. Here we call “LDpred2-low-h2” when running LDpred2(-grid) with a grid of hyper-parameters including these low values for *h*^2^. Results with LDpred2-low-h2 improves from 0.159 to 0.169 when using true sample sizes and from 0.134 to 0.163 when using the maximum sample size. Finally, we introduce a last change for robustness here: we form independent LD blocks in the LD matrix to prevent small errors in the Gibbs sampler to propagate to too many variants (Methods section 4.6). This change seems to solve convergence issues of LDpred2 in these simulations (Figure S3) and further improves predictive performance for all LDpred2 methods (Figure 1).

### 2.2 Misspecification when using imputed allele dosages

Marchini and Howie (2010) showed that the IMPUTE INFO measure is highly concordant with the MACH measure 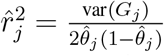, where 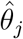 is the estimated allele frequency of *G* _*j*_, the genotypes for variant *j*. Therefore, when using the expected genotypes from imputation (allele dosages), their standard deviations are often lower than the expected value under Hardy–Weinberg equilibrium, because 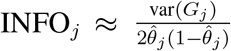. In simulations (cf. Methods section “Data for simulations”), we verify that sd 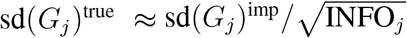 (Figure S5). As a direct consequence of the lower standard deviation of dosages, we also show that GWAS effect sizes 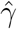 computed from imputed dosages are overestimated 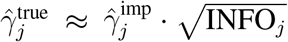 and 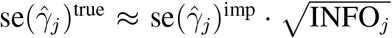 (Figures S6 and S7). This is the first correction of summary statistics we consider in the simulations below. As a second option, instead of using dosages to compute the GWAS summary statistics, it has been argued that using multiple imputation (MI) would be more appropriate (Palmer and Pe’er, 2016). In simulations, we show that 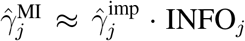 and 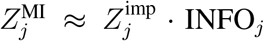, where 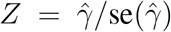 (Figure S8). This is the second correction of summary statistics we implement in the simulations below, along with *n*_*j*_ · INFO_*j*_ as the new per-variant sample sizes. Finally, we consider an in-between solution as a third correction, using 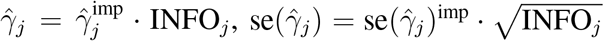 and *n*_*j*_ · INFO_*j*_ as the per-variant sample sizes. Note that we have recomputed INFO scores for the subset of 362,307 European individuals used in this paper since they can differ substantially from the ones reported by the UK Biobank for the whole data (Figures S9 and S10).

To compare these three corrections, we conduct a simulation using 40,000 variants from chromosome 22, where we simulate quantitative phenotypes assuming a heritability of 20% and 2000 causal variants using the “true” dataset (cf. Methods section “Data for simulations”). We compute GWAS summary statistics from the dosage dataset and use these summary statistics to run LDpred2 and lassosum2 with either no correction of the summary statistics, or with one of the three corrections described above. The LD reference used by LDpred2 and lassosum2 is computed from the validation set using the dataset with the “true” genotypes. For lassosum2 and LDpred2(-grid), which tune parameters using the validation set, correcting for imputation quality slightly improves predictive performance in these simulations (Figure 2). However, correcting for imputation quality can dramatically improve predictive performance for LDpred2-auto, provided the imputation quality is well estimated. Moreover, new additions for robustness introduced before, namely LDpred2-low-h2, LDpred2-auto-rob, and forming independent blocks in the LD matrix also improve predictive performance for all corrections (Figure 2).

**Figure 2:**
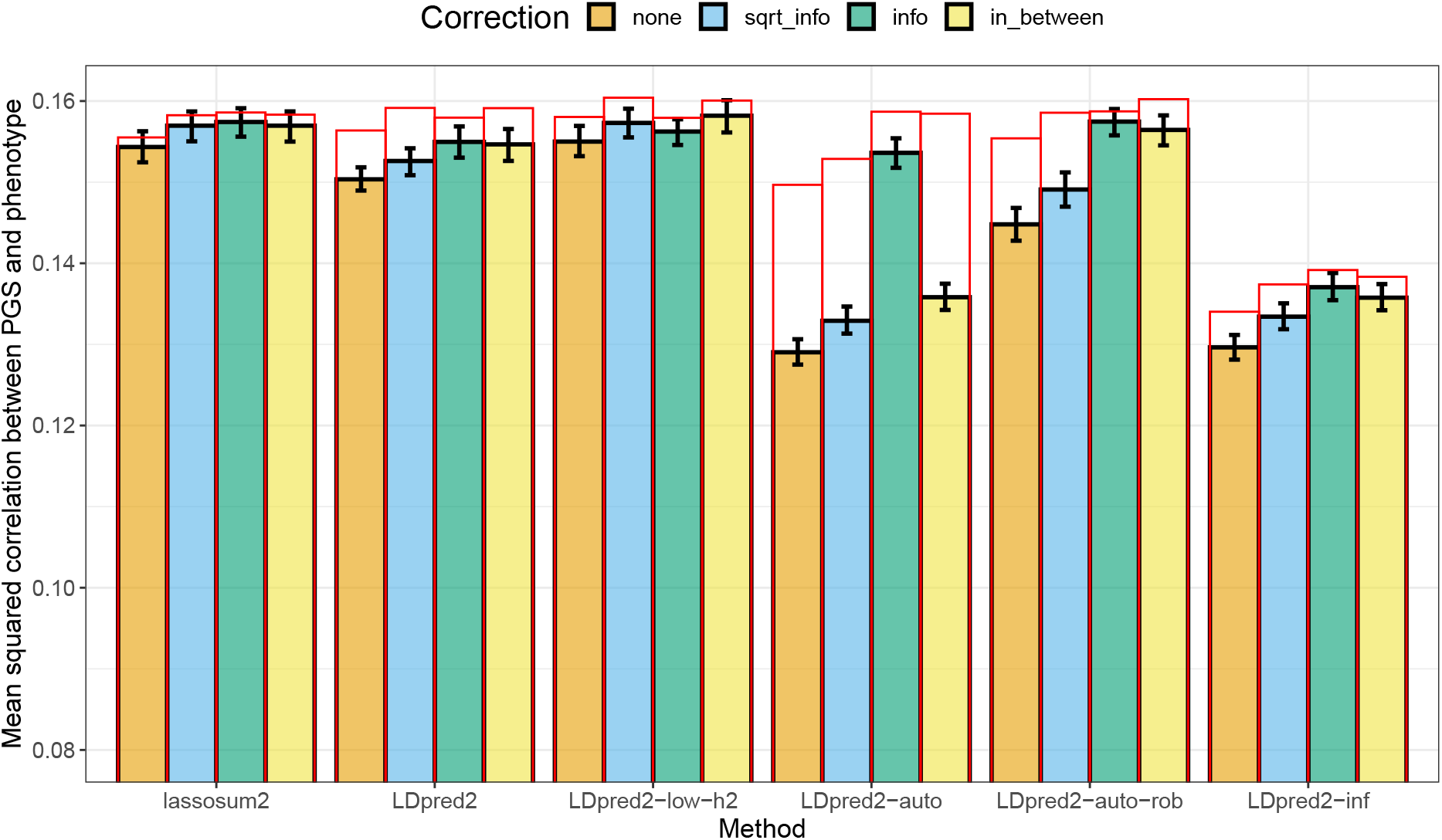
Results of predictive performance for the simulations using GWAS summary statistics from imputed dosage data, averaged over 10 simulations for each scenario. Reported 95% confidence intervals are computed from 10,000 non-parametric bootstrap replicates of the mean. Correction “sqrt_info” corresponds to using 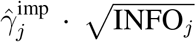 and 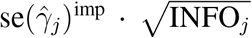. Correction “info” corresponds to using 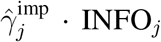 and *n*_*j*_ · INFO_*j*_. Correction “in_between” corresponds to using 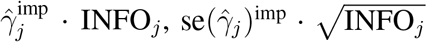 and *n*_*j*_ · INFO_*j*_. Red bars correspond to using the LD with independent blocks (Methods section 4.6).

In the real data applications hereinafter, we choose to use the first correction, “sqrt_info”, which is simple because it is equivalent to post-processing PGS effects by multiplying them by 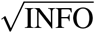.

### 2.3 Mismatch between LD reference and GWAS summary statistics

Here we design simulations to understand the impact of using a mismatched LD reference panel, e.g. that comes from a different population compared to the one used to compute the GWAS summary statistics. We use the same simulation setup as before (Methods). In addition, we design an alternative reference panel based on 10,000 individuals from South Europe by using the “Italy” center defined in Privé *et al*. (2022). When using this alternative LD reference panel instead of a well-matched one as in the previous sections, squared correlations between the polygenic scores and the simulated phenotypes drop from 0.174 to 0.167 for lassosum2, from 0.168 to 0.147 for LDpred2(-grid), from 0.174 to 0.169 for LDpred2-low-h2, from 0.143 to 0.139 for LDpred2-auto, from 0.162 to 0.146 for LDpred2-auto-rob, and from 0.144 to 0.140 for LDpred2-inf (Figure 3, averaging over 10 simulations). Forming independent LD blocks in these two LD matrices always helps, but do not improve performance by much when using the alternative LD.

**Figure 3:**
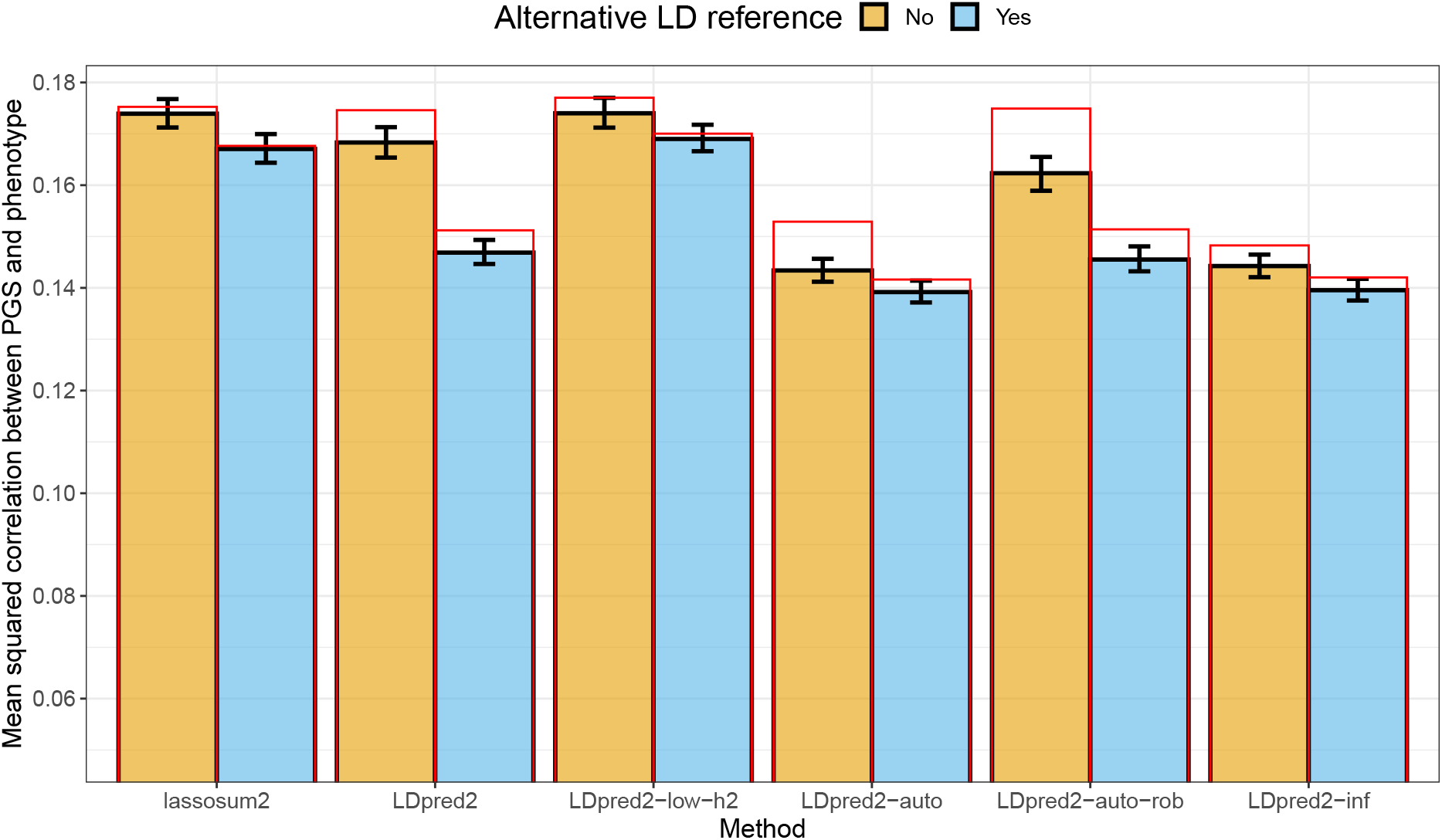
Results for the simulations with summary statistics with LD matrices based on two different populations. One comes from the same population used for computing the GWAS summary statistics (North-West Europe), while the other one comes from South Europe (alternative LD reference). Reported 95% confidence intervals are computed from 10,000 non-parametric bootstrap replicates of the mean. Red bars correspond to using the LD with independent blocks (Methods section 4.6).

### 2.4 Application to breast cancer summary statistics

In this section, we transition to using real data. Breast cancer GWAS summary statistics are interesting because they include results from two mega analyses (Michailidou *et al*., 2013, 2015, 2017), which means that parameters reported in these GWAS summary statistics, such as INFO scores and sample sizes, are estimated with high precision. Imputation INFO scores for the OncoArray summary statistics are generally very good (mean of 0.968 after having restricted to HapMap3 variants, Figure S13) and better than the ones from iCOGS (mean of 0.841, Figure S12), probably because the iCOGS chip included around 200K variants only, compared to more than 500K variants for the OncoArray. For both summary statistics, we compare the standard deviations (of genotypes) inferred from the reported allele frequencies (i.e. 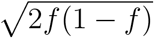 where *f* is the allele frequency, and denoted as sd_af) versus the ones inferred from the GWAS summary statistics (Equation (2), and denoted as sd_ss). As shown in figures 4 and S16, there is a clear trend with sd_ss being lower than sd_af as INFO decreases; indeed, using 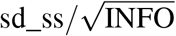 provides a very good fit for sd_af, except for some variants of chromosome 6 and 8 for the OncoArray summary statistics. Most of these outlier variants are either in region 25-33 Mbp of chromosome 6 or in 8-12 Mbp of chromosome 8 (Figure S14), which are two known long-range LD regions (Price *et al*., 2008). We hypothesize that this is due to using principal components (PCs) that capture LD structure instead of population structure, as covariates in the GWAS (Privé *et al*., 2020a). To validate this hypothesis, we simulate a phenotype using HapMap3 variants of chromosome 6 for 10,000 individuals from the UK Biobank, then we run GWAS with or without PC19 as covariate. PC19 from the UK Biobank was previously reported to capture LD structure in region 70-91 Mbp of chromosome 6 (Privé *et al*., 2020a). In these simulations, the same bias as in figure 4B is observed for the variants in this region (Figure S15), confirming our hypothesis.

**Figure 4:**
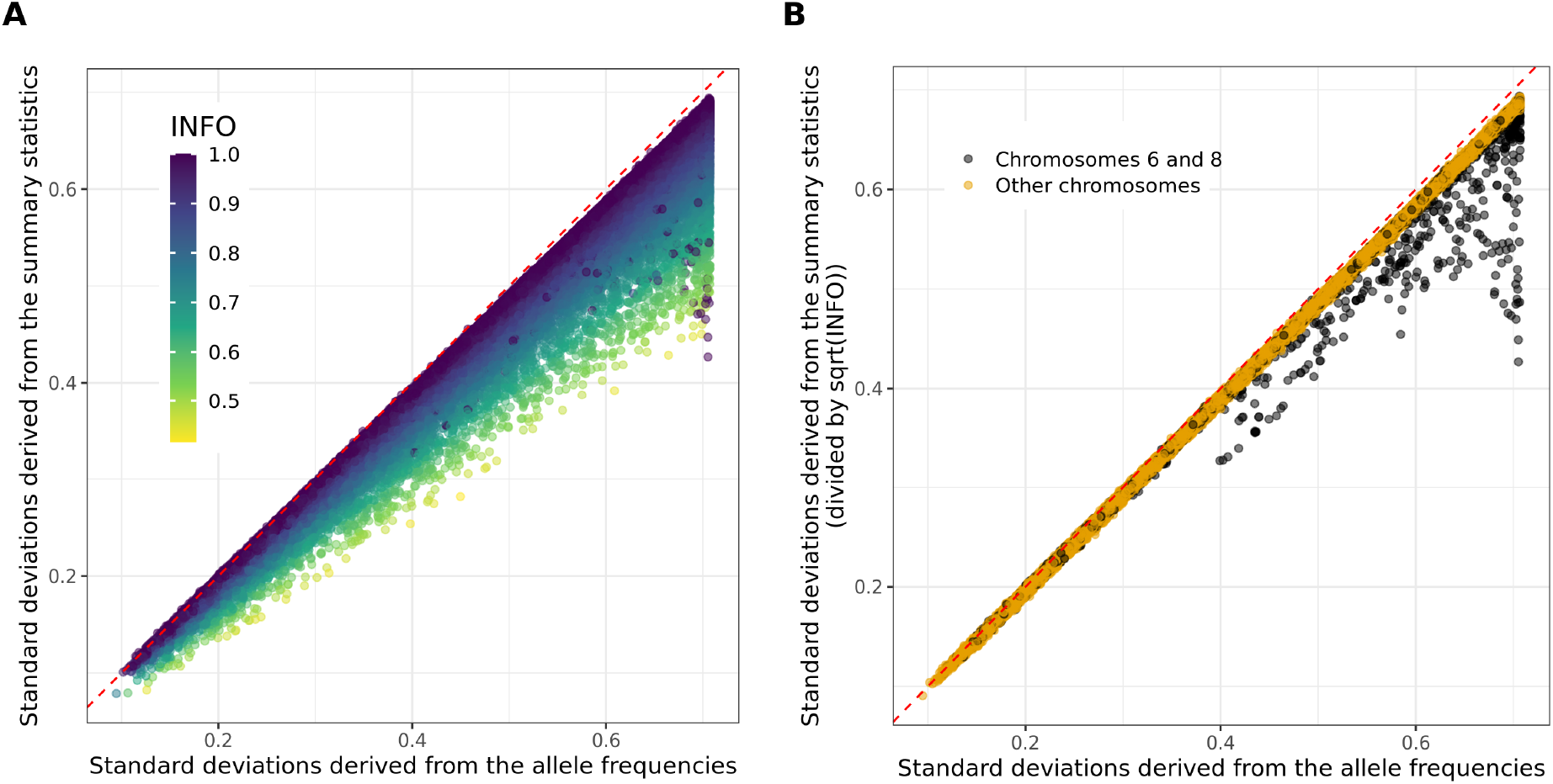
Standard deviations inferred from the OncoArray breast cancer GWAS summary statistics using equation (2) (**A:** Raw or **B:** dividing them by 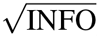) versus the ones inferred from the reported GWAS allele frequencies *f*_*j*_ (using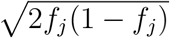). Only 100,000 HapMap3 variants are represented, at random.

Therefore, providing an accurate imputation INFO score is useful for two reasons. First, it allows for correcting for a reduced standard deviation when using imputed data in the QC step we propose, in order to better uncover possible problems with the GWAS summary statistics. Second, using one of the proposed corrections based on INFO scores may lead to an improved prediction when deriving polygenic scores. We apply these corrections to the two breast cancer summary statistics. In the comparison, we use the QC proposed in Privé et al. (2020b) first (which ends up filtering on MAF here, which we call “qc1”). We then also filter out the two long-range LD regions of chromosome 6 and 8 for the OncoArray summary statistics and remove around 500 variants when filtering on differences of AFs between summary statistics and the validation dataset (“qc2”). As for the correction using INFO scores, we use the first correction, “sqrt_info”, which is simple because it is equivalent to post-processing PGS effects by multiplying them By 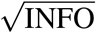. Although results for both QC used are very similar, correcting using INFO scores slightly improves predictive performance when deriving polygenic scores based on iCOGS summary statistics (Figure S22). All other improvements introduced before have little to no effect here, probably because misspecifications are much smaller than in the simulations.

### 2.5 Results for other phenotypes

We use other external GWAS summary statistics for which INFO scores are reported (see Table 2); they all have a very high mean INFO score (larger than 0.94), except for T1D-affy (0.885). QC plots comparing standard deviations usually show little deviation from the identity line (after the INFO score correction), except for coronary artery disease (CAD) summary statistics (Figures S16-S21). In figures 5 and S23-S27, we then provide similar results as figure S22 for other phenotypes. For MDD and PRCA, most changes introduced before have little to no impact on predictive performance. For CAD and T1D, the QC proposed in Privé *et al*. (2020b) and the additional QC proposed here provides much better predictive performance, especially for LDpred2-auto (and LDpred2-auto-rob) than when using no quality control, showing how important this preliminary step is.

**Table 2:** Summary of external GWAS summary statistics used. PRCA summary statistics have many variants with a missing INFO score, which we discard. We also restrict to variants with an INFO score larger than 0.4. Vitamin D summary statistics do not report INFO scores.

| Trait | GWAS citation | Effective GWAS sample size | # GWAS variants | # matched variants with INFO > 0.4 | Mean INFO |
| --- | --- | --- | --- | --- | --- |
| Breast cancer (BRCA) [iCOGS] | Michailidou *et al.* (2017) | 87,037 | 11,792,542 | 1,051,242 | 0.841 |
| Breast cancer (BRCA) [OncoArray] | Michailidou *et al.* (2017) | 104,442 | 11,792,542 | 1,054,233 | 0.968 |
| Type 1 diabetes (T1D) [Affymetrix] | Censin *et al.* (2017) | 5516 | 8,996,866 | 934,712 | 0.885 |
| Type 1 diabetes (T1D) [Illumina] | Censin *et al.* (2017) | 7982 | 8,996,866 | 949,334 | 0.942 |
| Prostate cancer (PRCA) | Schumacher *et al.* (2018) | 135,316 | 20,370,946 | 818,400 | 0.969 |
| Depression (MDD) [without UKBB] | Wray *et al.* (2018) | 110,464 | 9,874,289 | 1,049,455 | 0.968 |
| Coronary artery disease (CAD) | Nikpay *et al.* (2015) | 129,014 | 9,455,778 | 1,052,200 | 0.982 |
| Vitamin D | Jiang *et al.* (2018) | 79,366 | 2,579,296 | 1,016,935 | NA |

**Figure 5:**
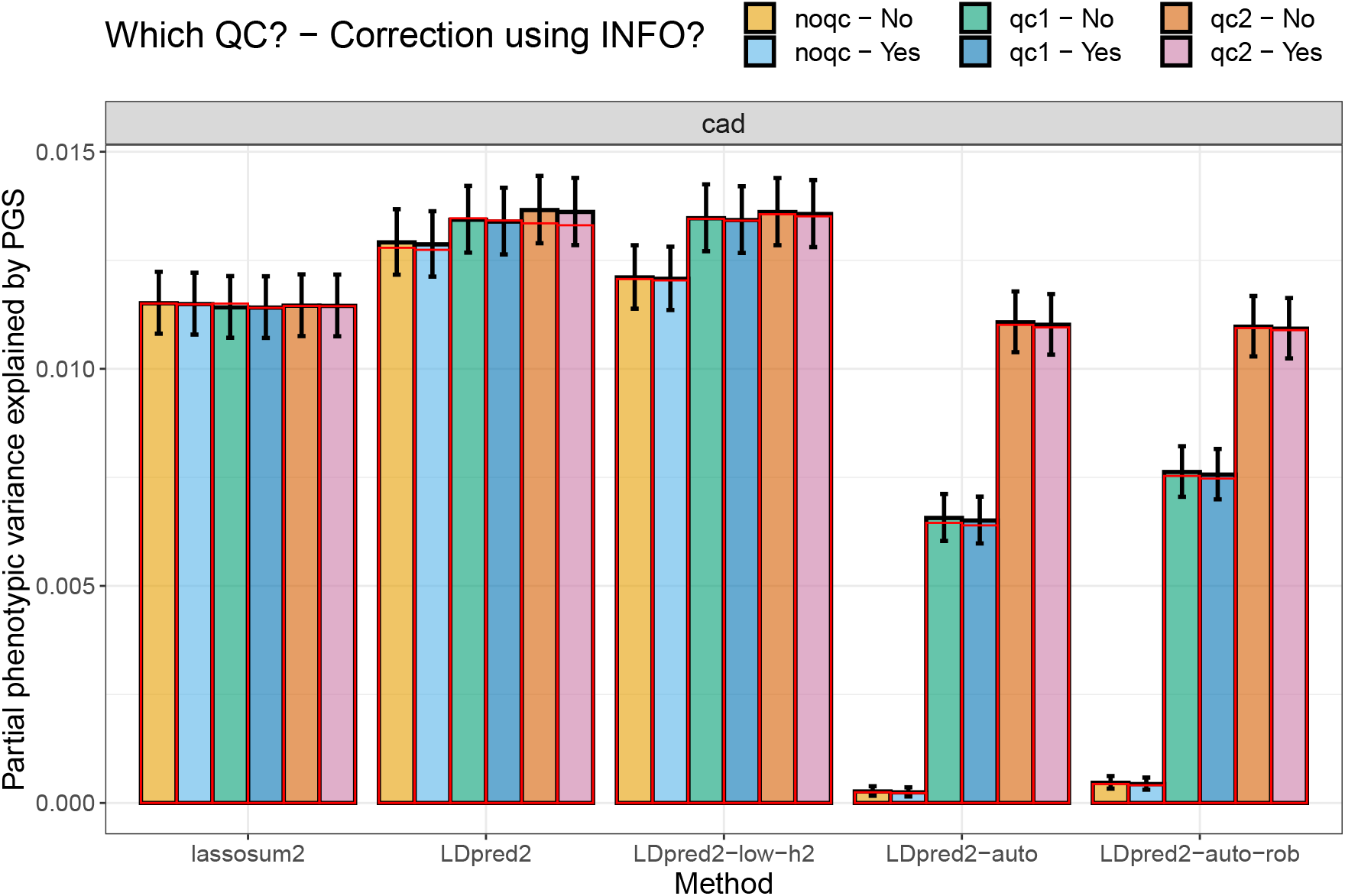
Variance explained of CAD in the UK Biobank by PGS derived from external summary statistics. These are computed using function `pcor` of R package bigstatsr where 95% confidence intervals are obtained through Fisher’s Z-transformation; these values are then squared. Red bars correspond to using the LD with independent blocks (Methods).

We also investigate results for vitamin D GWAS summary statistics, which do not report INFO scores nor allele frequencies, as opposed to previous ones. The QC procedure is then less precise and uses allele frequencies from the LD reference (Figure S28). However, these summary statistics do report per-variant sample sizes, which cover a wide range of different values (Figure S29). Here we compare using either the maximum sample size or the true per-variant sample sizes when deriving polygenic scores, and an additional QC step “qc2” which refers to removing all variants with a per-variant sample size less than 70% of the maximum one. When the true per-variant sample sizes are used, the additional “qc2” does not seem to be necessary (Figure 6), which is a bit surprising to us. When the maximum sample size is used, this “qc2” is very important, especially for LDpred2-auto. Note that, when using the maximum sample size to derive “sd_ss” in “qc1” (not the case here; we used the per-variant sample sizes), “qc1” would remove variants with underestimated standard deviations due to overestimated sample sizes, which would effectively remove variants with low sample sizes (Figure S1), similar to the “qc2” used here.

**Figure 6:**
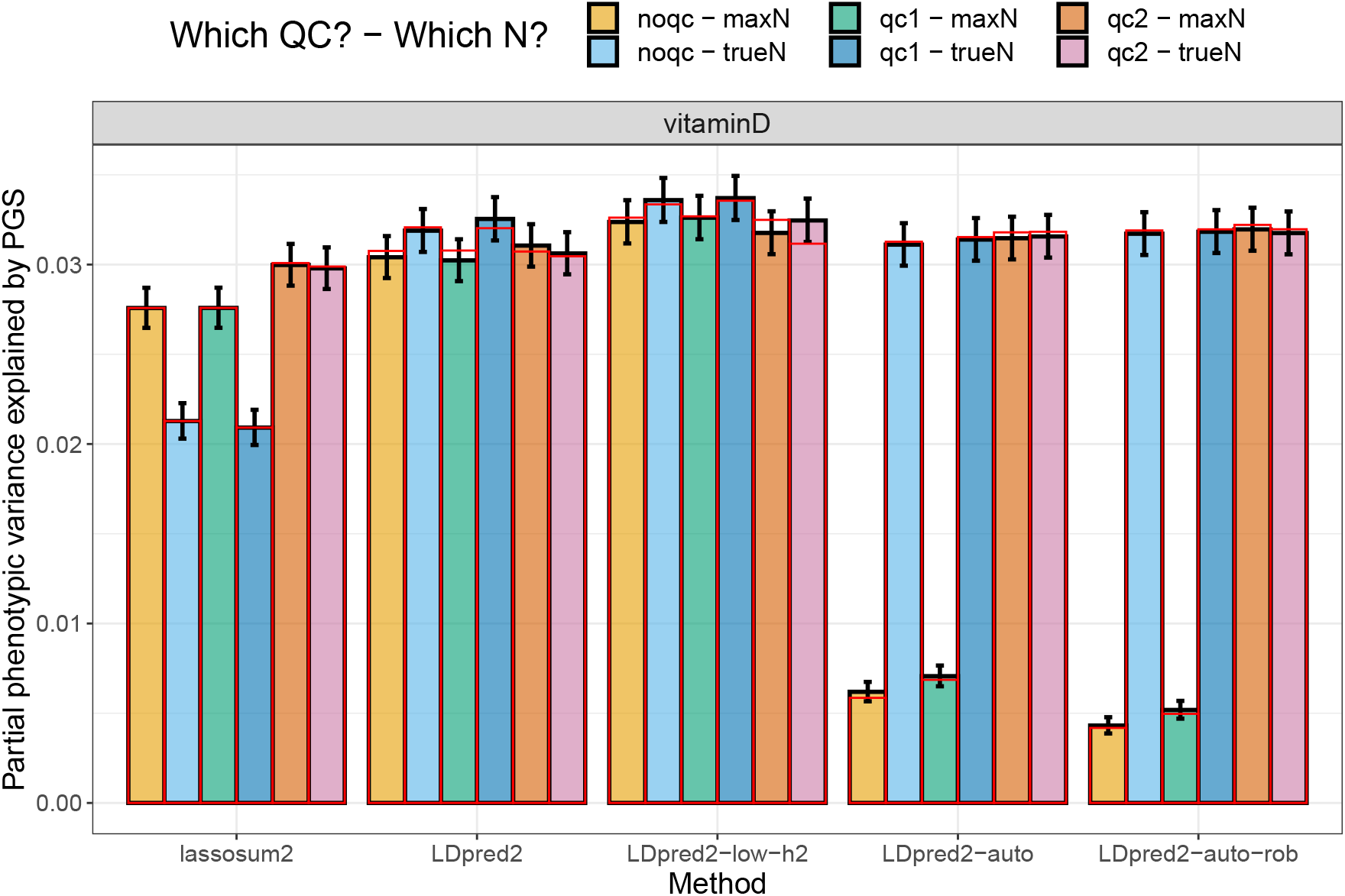
Variance explained of vitamin D in the UK Biobank by PGS derived from external summary statistics. These are computed using function `pcor` of R package bigstatsr where 95% confidence intervals are obtained through Fisher’s Z-transformation; these values are then squared. Red bars correspond to using the LD with independent blocks (Methods).

### 2.6 Application to FinnGen and Biobank Japan summary statistics

Here we investigate the use of different LD reference panels with GWAS summary statistics from two large biobanks of isolated populations. We use GWAS summary statistics for five disease endpoints from FinnGen (release 6, Kurki et al. (2022)), namely breast and prostate cancers (BrCa and PrCa), coronary artery disease (CAD), type 1 and type 2 diabetes (T1D and T2D). These were derived using SAIGE (Zhou *et al*., 2018). First, for each phenotype, we estimate a global effective sample using the 80th percentile of imputed sample sizes from equation (4). This estimation is only 84.7% for BrCa, 79.1% for PrCa, 73.1% for T1D, 64.5% for T2D, and 61.3% for CAD, when compared to the effective sample computed from the reported numbers of cases and controls. We believe this reduction in effective sample size is due to both having related individuals included in the analyses, as well as using SAIGE, as noted in the introduction. We then compare three different LD reference panels to use with these GWAS summary statistics (Methods section 4.7). Interestingly, using the small Finnish LD reference panel composed of 503 individuals only we have defined here seems to consistently provide more predictive polygenic scores than using the large UK one composed of 10,000 individuals or the widely used European subset of the 1000 Genomes (1000G) data (Figure 7). Note that the PGS here are validated in people of UK-like ancestry in the UK biobank to obtain sufficient sample sizes.

**Figure 7:**
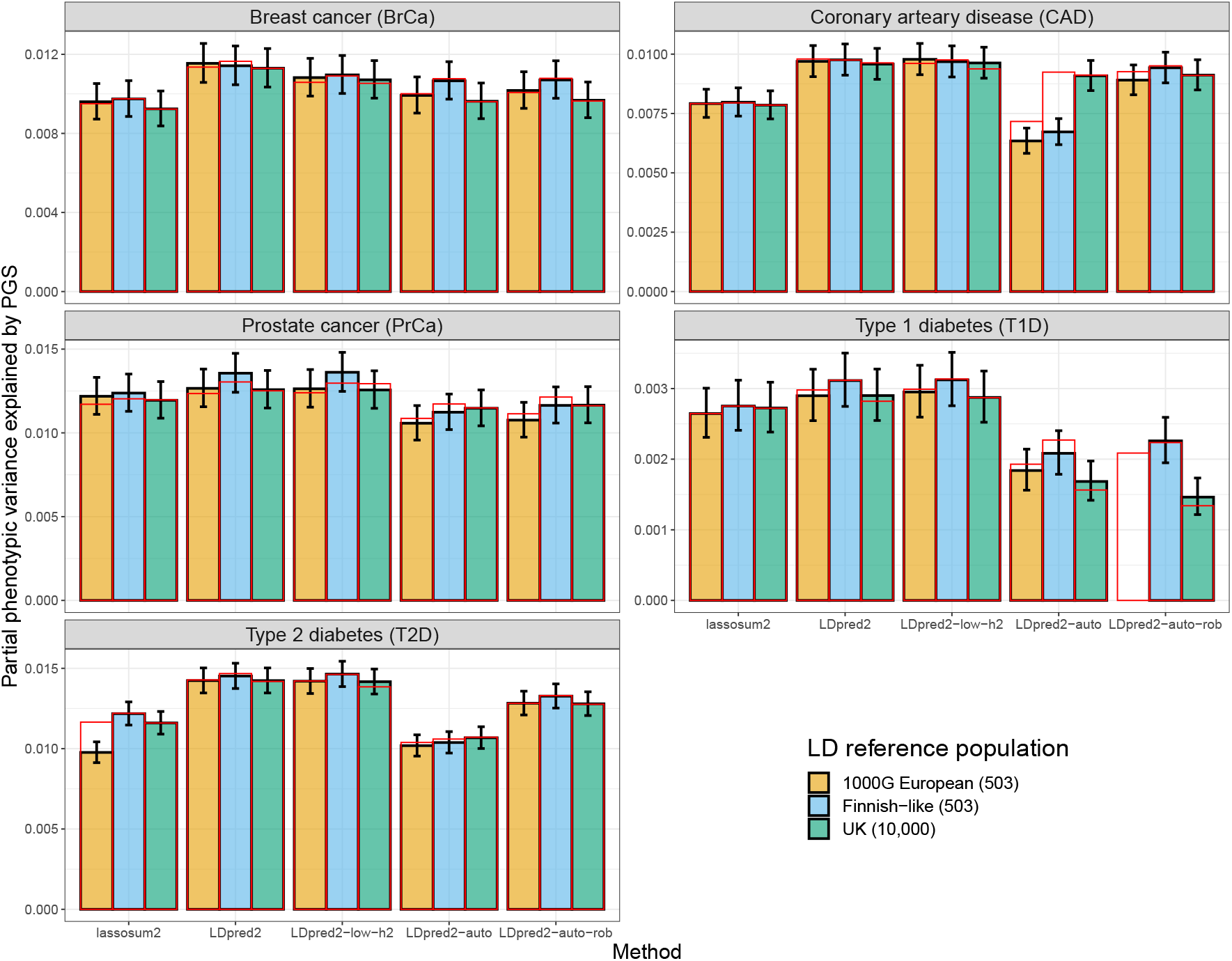
Results for PGS derived from five FinnGen GWAS summary statistics and using three different LD references. Partial correlations are computed using function `pcor` of R package bigstatsr where 95% confidence intervals are obtained through Fisher’s Z-transformation, then all values are squared to report the phenotypic variance explained by PGS. Red bars correspond to using the LD with independent blocks (Methods).

We also use GWAS summary statistics for four continuous outcomes from Biobank Japan (Sakaue *et al*., 2021). These were derived using BOLT-LMM-inf (Loh *et al*., 2018). First, for each phenotype, we estimate a global power improvement provided by BOLT-LMM by comparing *χ*^2^-statistics (the boost of BOLT-LMM-inf versus linear regression) across genome-wide significant variants, as recommended in Loh et al. (2018). This estimated power ratio, which corresponds to the increase in effective sample size, is 1.143 for height, 1.039 for HDL cholesterol, 1.025 for BMI, and 1.013 for systolic blood pressure. We then compare three different LD reference panels to use with these GWAS summary statistics (Methods section 4.7). Interestingly, when looking at height and to some extent at HDL cholesterol where we could get the best predictive performance, using the smaller Japanese LD reference we have defined here seems to provide more predictive PGS than using the one using a larger set of East Asian individuals from the UKBB or the widely used East Asian subset of the 1000 Genomes (1000G) data (Figure S30). Note that the PGS here are validated in people of broad East Asian ancestry in the UK biobank and for continuous outcomes to obtain sufficient sample sizes.

## 3 Discussion

Here we have investigated misspecifications in GWAS summary statistics, focusing particularly on the impact of sample size heterogeneity and imputation quality, and the application to polygenic scores (PGS) methods. Previously, we proposed a quality control (QC) based on comparing standard deviations (of genotypes) inferred from GWAS summary statistics with the ones computed from a reference panel (Privé *et al*., 2020b). Here we show that we can refine this QC by deriving the latter directly from the reported allele frequencies in the GWAS summary statistics, and by correcting the former using imputation INFO scores. Using this refined QC, we are able to identify a potential issue with how principal components were derived in a GWAS of breast cancer. Fortunately, this has practically no effect on the predictive performance of the derived polygenic scores. Additional QC can also be performed, e.g. comparing reported GWAS allele frequencies with the ones from the LD reference panel, e.g. to detect genotyping errors or allele inversions. We perform this additional QC as part of “qc2” here. One can also run other QC tools such as DENTIST (Chen *et al*., 2021), and also infer ancestry proportions from summary statistics to make sure these are matching with the LD reference used (Privé, 2021b).

Note that, in this study, we mostly use GWAS summary statistics that include extended information (e.g. INFO scores and allele frequencies), yet most GWAS summary statistics do not provide such exhaustive information (MacArthur *et al*., 2021). We acknowledge that, in the case of a meta-analysis from multiple studies, providing a single INFO score per variant may not be possible. Solutions such as using a weighted averaged INFO score might be worth exploring in future studies. Nevertheless, this quality control could be performed within each study before meta-analyzing results, to make sure that resulting summary statistics have the best possible quality for follow-up analyses such as deriving polygenic scores. Another information, the effective sample size per variant, is often missing from GWAS summary statistics. Sometimes, it can even be challenging to recover the total effective sample size from large meta-analyses. We recall that, when some studies have an imbalanced number of cases and controls, the total effective sample size of their meta-analysis should not be computed from the total numbers of cases and controls overall, but instead from the sum of the effective sample sizes of each study (Grotzinger *et al*., 2021). Indeed, take the extreme example of meta-analyzing two studies, one with 1000 cases and 0 controls, and another one with 0 cases and 1000 controls, then the effective sample size of the meta-analysis is 0, not 2000. Misspecifying the GWAS sample size can lead to serious issues such as misestimating the SNP heritability (Grotzinger *et al*., 2021). Fortunately, an overestimated sample size can be detected from the QC plot we propose, where the slope is then less than 1 for case-control studies using logistic regression; otherwise the standard deviation of the phenotype is also needed (Equation (1)), but can be estimated (Privé *et al*., 2020b). Here we have used this strategy to estimate reduced effective sample sizes in FinnGen GWAS summary statistics.

We have assessed the impact of these misspecifications in GWAS summary statistics on the predictive performance of some polygenic score methods. Using both the Bayesian LDpred2 models (Privé *et al*., 2020b) and our reimplementation of the frequentist lassosum model (Mak *et al*., 2017) for deriving polygenic scores, we have introduced and investigated some changes to possibly make these models more robust to misspecifications. Overall, these changes provided large improvements of predictive performance in the simulations with large misspecifications. The proposed quality controls on GWAS summary statistics also provided better predictive performance for CAD, T1D, and vitamin D. However, these changes had limited effect when applied to some other real GWAS summary statistics, which is both unfortunate but also reassuring because it means that these GWAS summary statistics are of particularly good quality for follow-up analyses such as deriving polygenic scores.

In conclusion, we recommend adopting these changes, i.e. performing the (refined) QC proposed here, forming independent LD blocks in the LD matrix, and using more regularization when needed. We also recommend using well-matched LD reference panels. More regularization can be achieved by testing additional smaller values for the heritability parameter in LDpred2-grid, and using the two new parameters introduced here in LDpred2-auto (Methods section 4.5). Note that LD blocks are already widely used by several methods, such as lassosum and PRS-CS, because they allow for processing smaller matrices at once (Mak *et al*., 2017; Ge et al., 2019). Imposing independent blocks on the LD matrix could result in further misspecifications (Zhou and Zhao, 2021), but here we have shown that well defined blocks can actually make PGS methods more robust. PRS-CS is currently one of the most robust PGS methods; for example, it can use the (small) 1000 Genomes data as LD reference (Ge *et al*., 2019), and can even use a European LD reference panel with multi-ancestry GWAS summary statistics (Wang *et al*., 2021). We believe this is made possible by the use of a strong regularization in PRS-CS (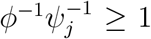, which would approximately correspond to using *s* = 0.5 in lassosum, *δ* = 1 in lassosum2, and the new parameter `shrink_corr` = 0.5 in LDpred2-auto). Using enough regularization is good for robustness, but note that using too much regularization can also damage predictive performance. To address this limitation, we recommend to rely on a proper QC and choice of ancestry-matched LD instead of on too much regularization. We would like to encourage large biobanks, such as FinnGen and Biobank Japan, to provide LD reference matrices matching the large GWAS summary statistics they provide, ideally based on the same large number of individuals. As a future research direction, we are interested in using multiancestry-matched LD matrices to use with multi-ancestry GWAS summary statistics in order to improve polygenic prediction in all ancestries. It would also be useful to investigate the impact of the QC, well-matched LD reference panels and other adjustments we propose here on other (non-PGS) methods, for which consistency and robustness are likely to be very important as well (Chen *et al*., 2021).

## 4 Materials and Methods

### 4.1 Data for simulations

We use the UK Biobank imputed (BGEN) data (Bycroft *et al*., 2018). We restrict individuals to the ones used for computing the principal components (PCs) in the UK Biobank (Field 22020). These individuals are unrelated and have passed some quality control including removing samples with a missing rate on autosomes larger than 0.02, having a mismatch between inferred sex and self-reported sex, and outliers based on heterozygosity (more details can be found in section S3 of Bycroft et al. (2018)). To get a set of genetically homogeneous individuals, we compute a robust Mahalanobis distance based on the first 16 PCs and further restrict individuals to those within a log-distance of 5 (Privé *et al*., 2020a). This results in 362,307 individuals. We randomly sample 300,000 individuals to form a training set (e.g. to run the GWAS), 10,000 individuals to form a validation set (to tune hyper-parameters), and use the remaining 52,307 individuals to form a test set (to evaluate final predictive models).

Among genetic variants on chromosome 22 and with a minor allele frequency larger than 0.01 and an imputation INFO score larger than 0.4 (as reported by the UK Biobank), we sample 40,000 of them according to the inverse of the INFO score density so that they have varying levels of imputation accuracy (Figure S4). We read the UK Biobank data into two different datasets using function `snp_readBGEN` from R package bigsnpr (Privé *et al*., 2018), one by reading the BGEN data at random according to imputation probabilities, and another one reading it as dosages (i.e. expected values according to imputation probabilities). The first dataset is used as what could be the real genotype calls and the second dataset as what would be its imputed version; this design technique was used in Privé et al. (2019).

### 4.2 Data for real analyses

We also use the UK Biobank data for validation/testing in real data analyses, and use the same individuals as described in the previous section. We sample 10,000 individuals to form a validation set and use the remaining 352,307 individuals as test set. We restrict to the 1,054,315 HapMap3 variants used in the LD reference provided in Privé et al. (2020b).

To define phenotypes in the UK Biobank, we first map ICD10 and ICD9 codes (UKBB fields 40001, 40002, 40006, 40013, 41202, 41270 and 41271) to phecodes using R package PheWAS (Carroll *et al*., 2014; Wu et al., 2019). We also use some continuous phenotypes, namely vitamin D (Data-Field 30890), height (50), BMI (21001), systolic blood pressure (4080), and HDL cholesterol (30760).

We use published GWAS summary statistics listed in table 2 to derive polygenic scores. We also use GWAS summary statistics for five disease endpoints from FinnGen (release 6, Kurki et al. (2022)) and for four continuous outcomes from Biobank Japan (Sakaue *et al*., 2021).

### 4.3 GWAS sample size imputation

In this paper, we extensively use the following formula

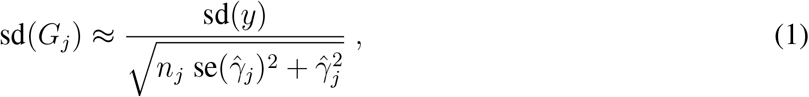

where 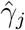 is the marginal GWAS effect size of variant *j, n*_*j*_ is the GWAS sample size associated with variant *j, y* is the vector of phenotypes and *G*_*j*_ is the vector of genotypes for variant *j*. This formula is used in LDpred2 Privé et al. (2020b, 2022). Note that, for a binary trait for which logistic regression is used, we have instead

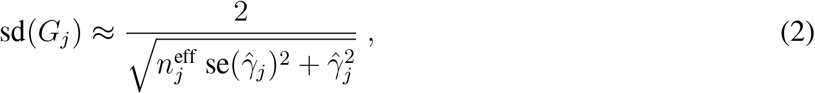

where 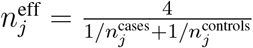.

We can then impute *n*_*j*_ from equation (1) using

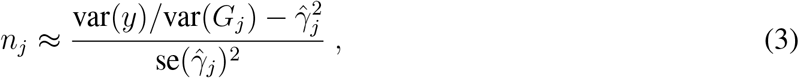

and impute 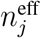 from equation (2) using

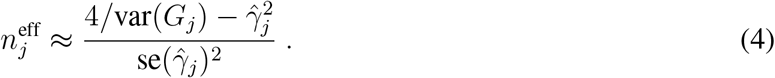

In practice, we also bound this estimate to be between 0.5 · *N* and 1.1 · *N*, where *N* is the total (effective) sample size. Also note that the estimate of var(*G*_*j*_) has to account for the imputation accuracy, i.e. use 2 · *f*_*j*_ · (1 − *f*_*j*_) ·INFO_*j*_ (Section 2.2).

### 4.4 New implementation of lassosum in bigsnpr

Instead of using a regularized version of the correlation matrix *R* parameterized by *s, R*_*s*_ = (1 −*s*)*R* +*sI* (where 0 *< s* ≤ 1), we use *R*_*δ*_ = *R* + *δI* (where *δ >* 0), which makes it clearer that lassosum is also using L2-regularization (therefore elastic-net). Then, from Mak et al. (2017), the solution from lassosum can be obtained by iteratively updating

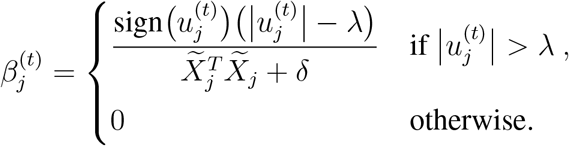

Where

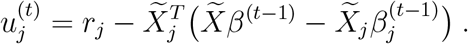

Following the notations from Privé et al. (2020b), denote 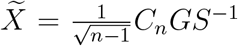, where *G* is the genotype matrix, *C*_*n*_ is the centering matrix and *S* is the diagonal matrix of standard deviations of the columns of *G*. Then 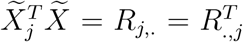 and 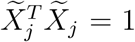. Moreover, using the notations from Privé et al. (2020b), 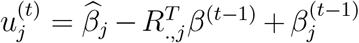, where 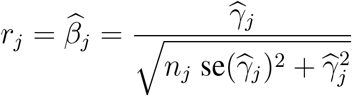 and 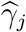 is the GWAS effect of variant *j* and *n* is the GWAS sample size (Mak *et al*., 2017; Privé et al., 2022). Then computing 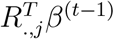 is the most time-consuming part of each iteration. To make this faster, instead of computing 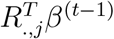 at each iteration (*j* and *t*), we can start with an initial vector of 0s only (for all *j*) since *β*^(0)^ ≡ 0, then update this vector when 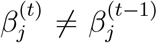 only. Note that only positions *k* for which *R*_*k,j*_ ≠ 0 must be updated in this vector 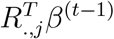.

In this new implementation of the lassosum model, which we call lassosum2, the input parameters are the correlation matrix *R*, the GWAS summary statistics (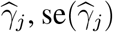 and *n*), and the two hyper-parameters *λ* and *δ*. Therefore, except for the two hyper-parameters, lassosum2 uses the exact same input parameters as LDpred2 (Privé *et al*., 2020b). We try *δ* ∈ {0.001, 0.005, 0.02, 0.1, 0.6, 3} by default in lassosum2, instead of *s* ∈ {0.2, 0.5, 0.8, 1.0} in lassosum. For *λ*, the default in lassosum uses a sequence of 20 values equally spaced on a log scale between 0.1 and 0.001. By default in lassosum2, we use a similar sequence, but between *λ*_0_ and *λ*_0_*/*100 instead, where 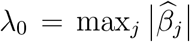 is the minimum *λ* for which no variable enters the model because the L1-regularization is too strong. Note that we do not provide an “auto” version using pseudo-validation (as in Mak et al. (2017)) as we have not found it to be robust enough (Figure S31). Also note that, as in LDpred2, we run lassosum2 genome-wide using a sparse correlation matrix which assumes that variants further away than 3 cM are not correlated. Although we do not require splitting the genome into independent LD blocks anymore (as done in lassosum), we recommend to do so for robustness in LDpred2, and for extra speed gain (cf. Methods section 4.6).

### 4.5 LDpred2-low-h2 and LDpred2-auto-rob

Here we introduce the small changes made to LDpred2 (-grid and -auto) in order to make them more robust. First, LDpred2-low-h2 simply consists in running LDpred2-grid while testing *h*^2^ within {0.3, 0.7, 1, 1.4}·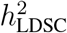 (note the added 0.3 compared to Privé et al. (2020b)), where 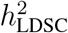 is the heritability estimate from LD score regression. Indeed, we show in simulations here that using lower values for *h*^2^ may provide higher predictive performance in the case of misspecifications (thanks to more shrinkage of the effects). In simulations, because of the large misspecifications, we use a larger grid over 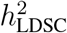.

For LDpred2-auto, we introduce two new parameters. The first one, `shrink_corr`, allows for shrinking off-diagonal elements of the correlation matrix. This is similar to parameter ‘s’ in lassosum (Methods section 4.4), and acts as a means of regularization. We use a value of 0.9 in simulations and 0.95 in real data when running “LDpred2-auto-rob” here, and the default value of 1 (with no effect) when running “LDpred2-auto”. The second new parameter, `allow_jump_sign`, controls whether variant effect sizes can change sign over two consecutive iterations of the Gibbs sampler. When setting this parameter to false (in the method we refer to as “LDpred2-auto-rob” here), this forces the effects to go through 0 before changing sign. This is useful to prevent instability (oscillation and ultimately divergence) of the Gibbs sampler under large misspecifications, and is also useful for accelerating convergence of chains with a large initial value for *p* (the proportion of causal variants).

### 4.6 New LD reference

We form nearly independent LD blocks using the optimal algorithm developed in Privé (2021a). For different numbers of blocks and maximum number of variants in each block, we use the split with the minimum cost within the ones reducing the original number of non-zero values to less than 60% (70% for chromosome 6). Having a correlation matrix with independent blocks prevents small errors in the algorithm (e.g. the Gibbs sampler in LDpred2) from propagating to too many variants. It also makes running LDpred2 (and lassosum2) faster, taking about 60% of the initial time (since only 60% of the initial non-zero values of the correlation matrix are kept).

We have also developed a new “compact” format for the SFBMs (sparse matrices on disk). Instead of using something similar to the standard “compressed sparse column” format which stores all {*i, x*(*i, j*)} for a given column *j*, we only store the first index *i*_0_ and all the contiguous values {*x*(*i*_0_, *j*), *x*(*i*_0_ + 1, *j*), … } up to the last non-zero value for this column *j*. This makes this format about twice as efficient for both LDpred2 and lassosum2 (in terms of both memory and speed). Therefore, using both this new format and the LD blocks should divide computation times by 3.

### 4.7 Alternative LD reference for FinnGen and Biobank Japan

To be used with GWAS summary statistics from FinnGen, we investigate three different LD reference panels. To define homogeneous ancestry groups in the UK Biobank, we include all individuals within a specific distance to a population center in the PCA space (Privé *et al*., 2022; Privé, 2021b). We use either the 503 European (including 99 Finnish) individuals from the 1000 Genomes (1000G) data (1000 Genomes Project Consortium *et al*., 2015), 404 Finnish-like UKBB + the 99 Finnish 1000G individuals (also 503 in total), or the 10,000 UK individuals from the validation set we use in this paper.

To be used with GWAS summary statistics from Biobank Japan, we also investigate three different LD reference panels. We use either the 504 East Asian (including 104 Japanese) individuals from the 1000G, 400 Japanese-like UKBB + the 104 Japanese 1000G individuals (also 504 in total), or 2041 East Asian UKBB individuals (including the 400 previous Japanese individuals). We also construct versions of these LD references with independent LD blocks, as described in the previous section.

## Code and data availability

All code used for this paper is available at https://github.com/privefl/paper-misspec/tree/master/code. We have extensively used R packages bigstatsr and bigsnpr (Privé *et al*., 2018) for analyzing large genetic data, packages from the future framework (Bengtsson, 2021) for easy scheduling and parallelization of analyses on the HPC cluster, and packages from the tidyverse suite (Wickham *et al*., 2019) for shaping and visualizing results.

The latest version of R package bigsnpr can be installed from GitHub, and a recent enough version can be installed from CRAN. A tutorial on running LDpred2 and lassosum2 using R package bigsnpr is available at https://privefl.github.io/bigsnpr/articles/LDpred2.html.

We have recomputed the allele frequencies and imputation INFO scores within the European subset used here and across all imputed variants in the UK Biobank, and made them available at https://doi.org/10.6084/m9.figshare.16635388. We have formed independent LD blocks within the Northwestern European LD references provided in Privé et al. (2020b), and made these updated versions available at https://doi.org/10.6084/m9.figshare.19213299.

## Acknowledgements

Authors thank Timothy Shin Heng Mak, Shing Wan Choi, and Matthew Stephens for helpful discussions. Authors also thank GenomeDK and Aarhus University for providing computational resources and support that contributed to these research results. This research has been conducted using the UK Biobank Resource under Application Number 58024.

