## Supplementary Materials for "Identifying and correcting for misspecifications in GWAS summary statistics and polygenic scores"

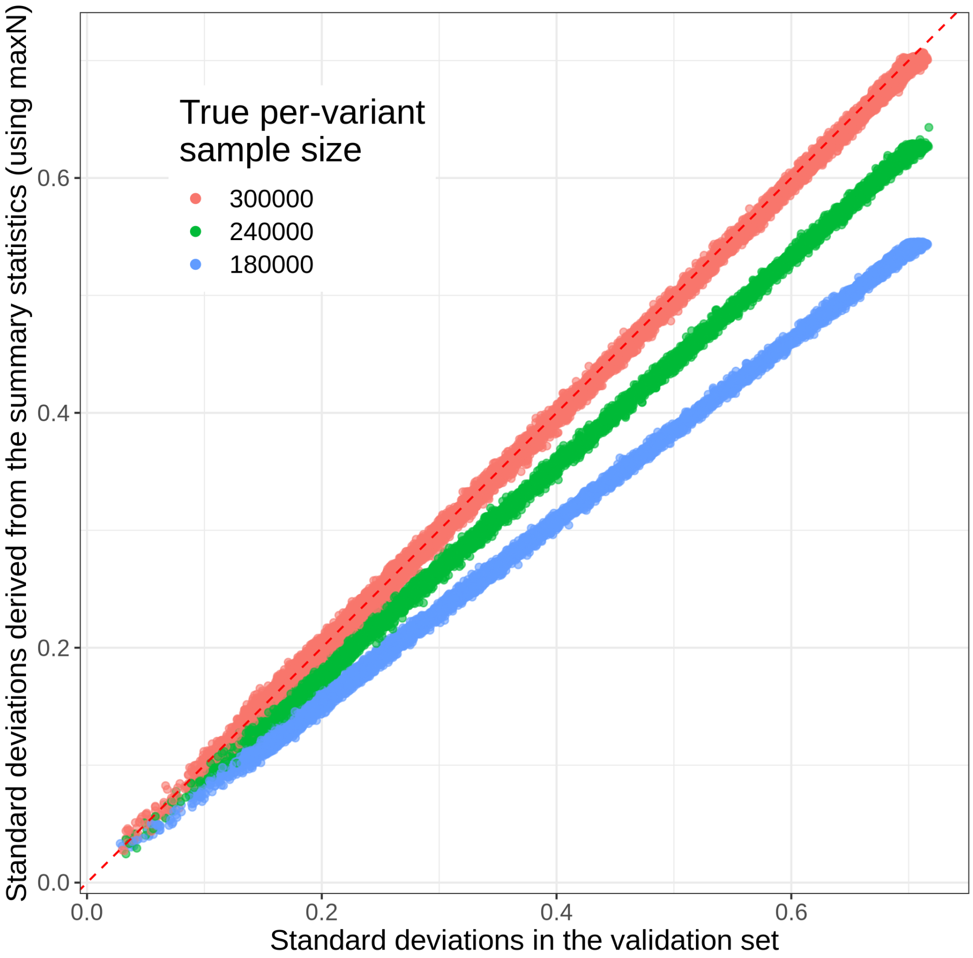


Figure S1: Quality control plot, as proposed in Privé *et al.* (2020), for the simulations with sample size misspecification. The standard deviations are derived from the summary statistics assuming the same global GWAS sample size for all variants (300,000).

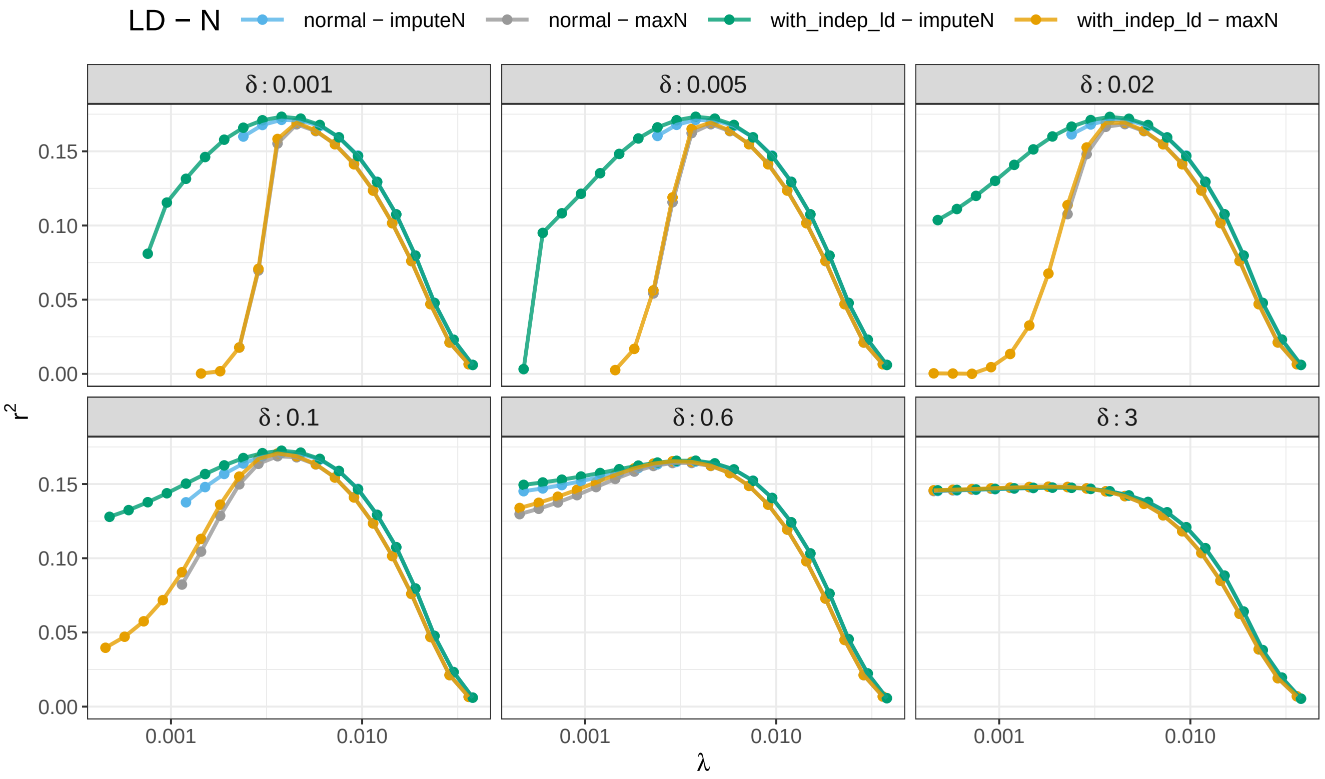


Figure S2: Multiple results (squared correlation $r^2$ between polygenic score and phenotype) over a grid of parameters ($\lambda$ and $\delta$) for lassosum2 in one of the simulations with misspecified GWAS sample sizes ($n_j$). The different colors indicate whether we use the maximum of $n_j$'s ("maxN") or the imputed $n_j$'s ("imputeN"), and the normal LD matrix ("normal") or the one with independent LD blocks ("with_indep_ld"). Missing points represent models that were detected as divergent.

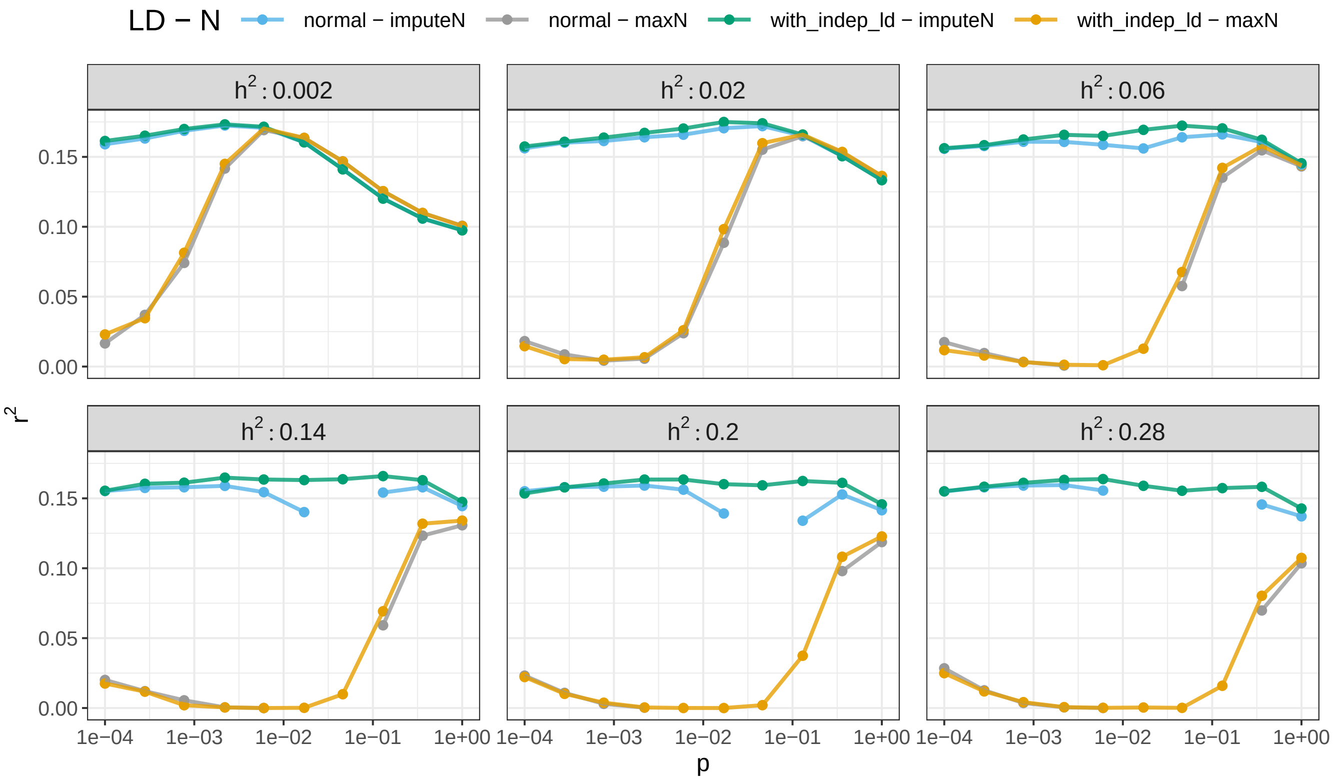


Figure S3: Multiple results (squared correlation $r^2$ between polygenic score and phenotype) over a grid of parameters (the SNP heritability $h^2$ and the proportion of causal variants $p$) for LDpred2(-grid) in one of the simulations with misspecified GWAS sample sizes ($n_j$). The different colors indicate whether we use the maximum of $n_j$'s ("maxN") or the imputed $n_j$'s ("imputeN"), and the normal LD matrix ("normal") or the one with independent LD blocks ("with_indep_ld"). Note that the small $h^2$ values (0.002, 0.02, 0.06) were added and are part of the method we call "LDpred2-low-h2" in the main text. Missing points represent models that were detected as divergent.

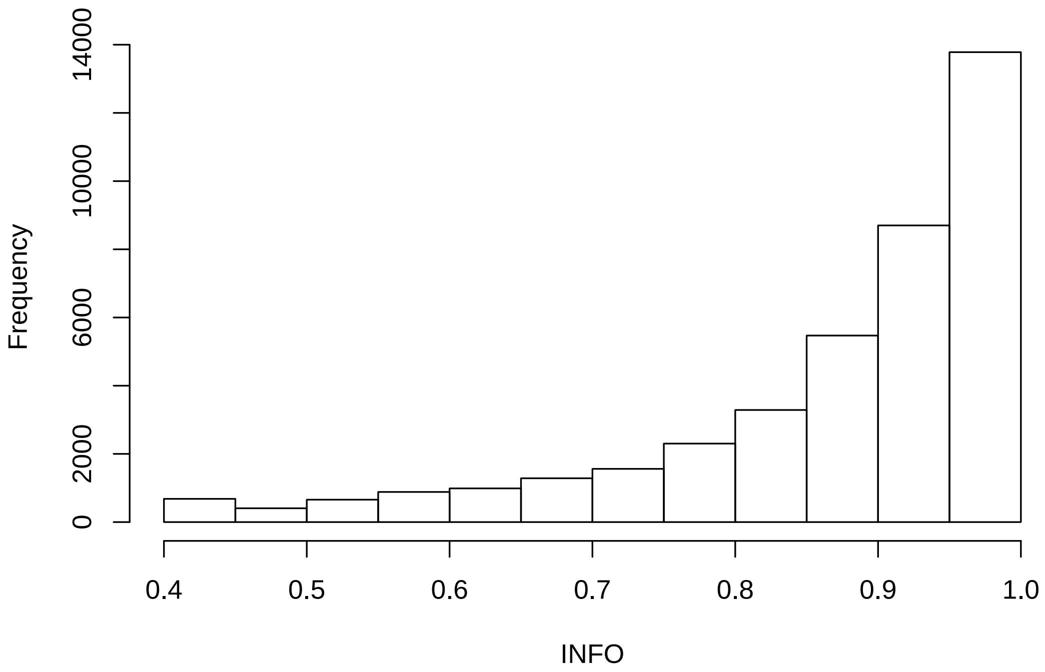


Figure S4: Histogram of the imputation INFO scores of the 40,000 variants from chromosome 22 of the UK Biobank data used in the simulations.

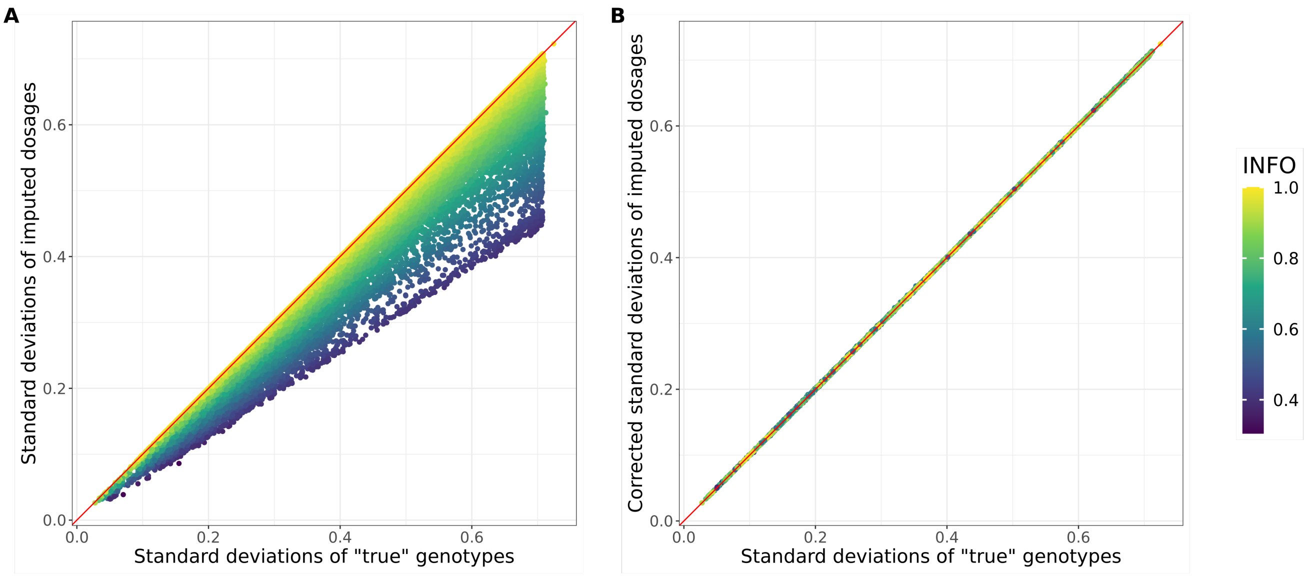


Figure S5: **A:** Raw standard deviations (SDs) from imputed dosages or **B:** corrected SDs from imputed dosages (dividing them by $\sqrt{\text{INFO}}$) versus SDs from "true" genotype calls, colored by INFO scores.

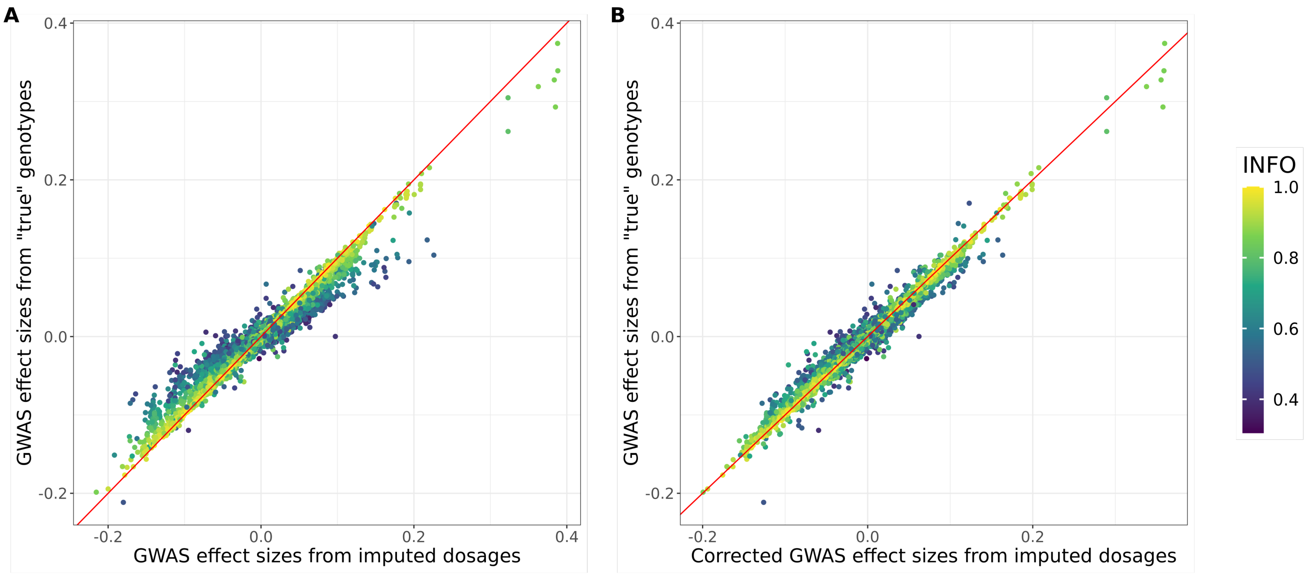


Figure S6: GWAS effect sizes ($\hat{\gamma}$) from "true" genotype calls versus either **A:** raw $\hat{\gamma}$ from imputed dosages or **B:** corrected $\hat{\gamma}$ from imputed dosages (multiplying them by $\sqrt{\text{INFO}}$), colored by INFO scores.

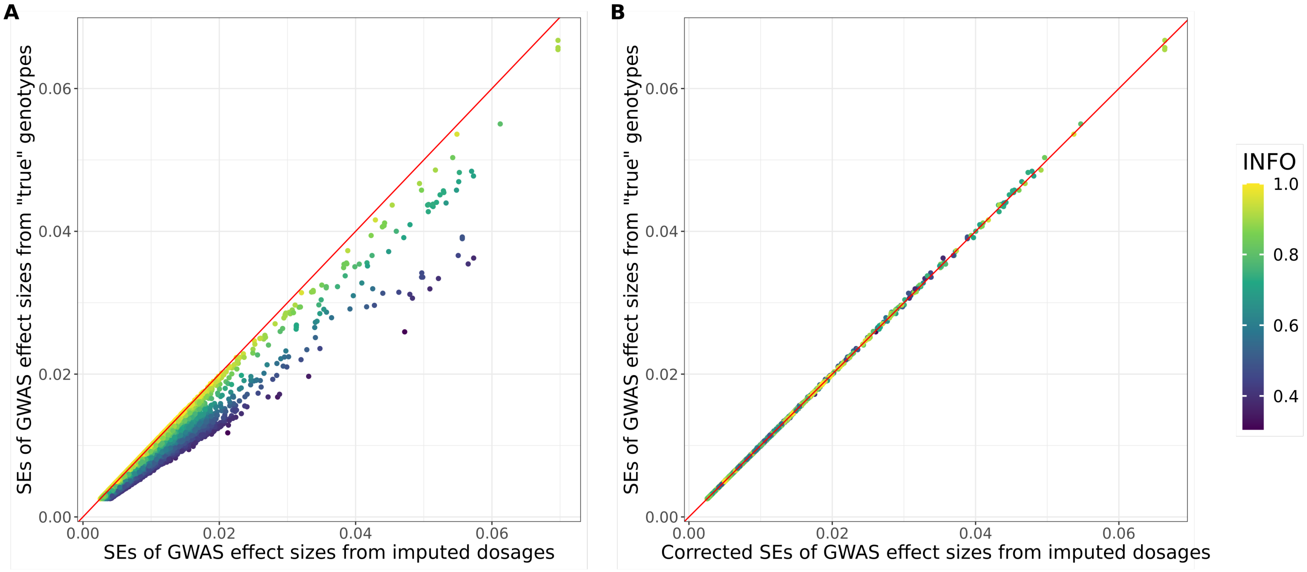


Figure S7: Standard errors of GWAS effect sizes (SEs) from “true” genotype calls versus either **A:** raw SEs from imputed dosages or **B:** corrected SEs from imputed dosages (multiplying them by $\sqrt{\text{INFO}}$), colored by INFO scores.

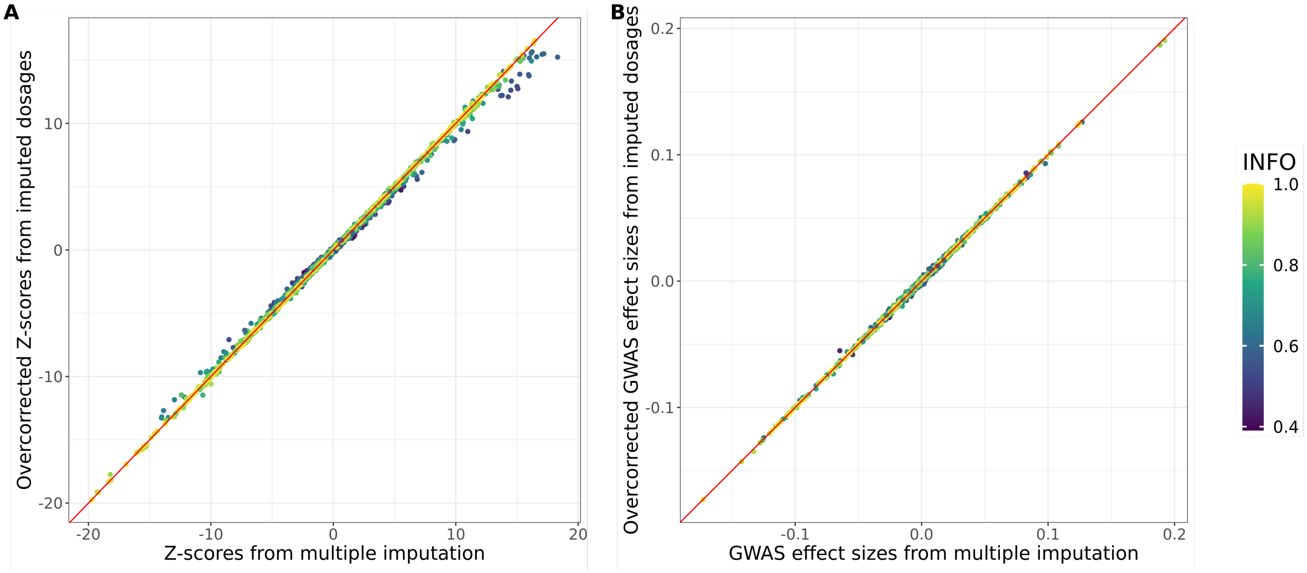


Figure S8: (Over)corrected GWAS **A:** Z-scores and **B:** effect sizes from imputed dosages (multiplying them by INFO) versus the ones obtained from multiple imputation (MI, 20 draws used, Palmer and Pe’er (2016)), colored by INFO scores. This shows only one tenth of the simulated variants, because MI can be computationally intensive.

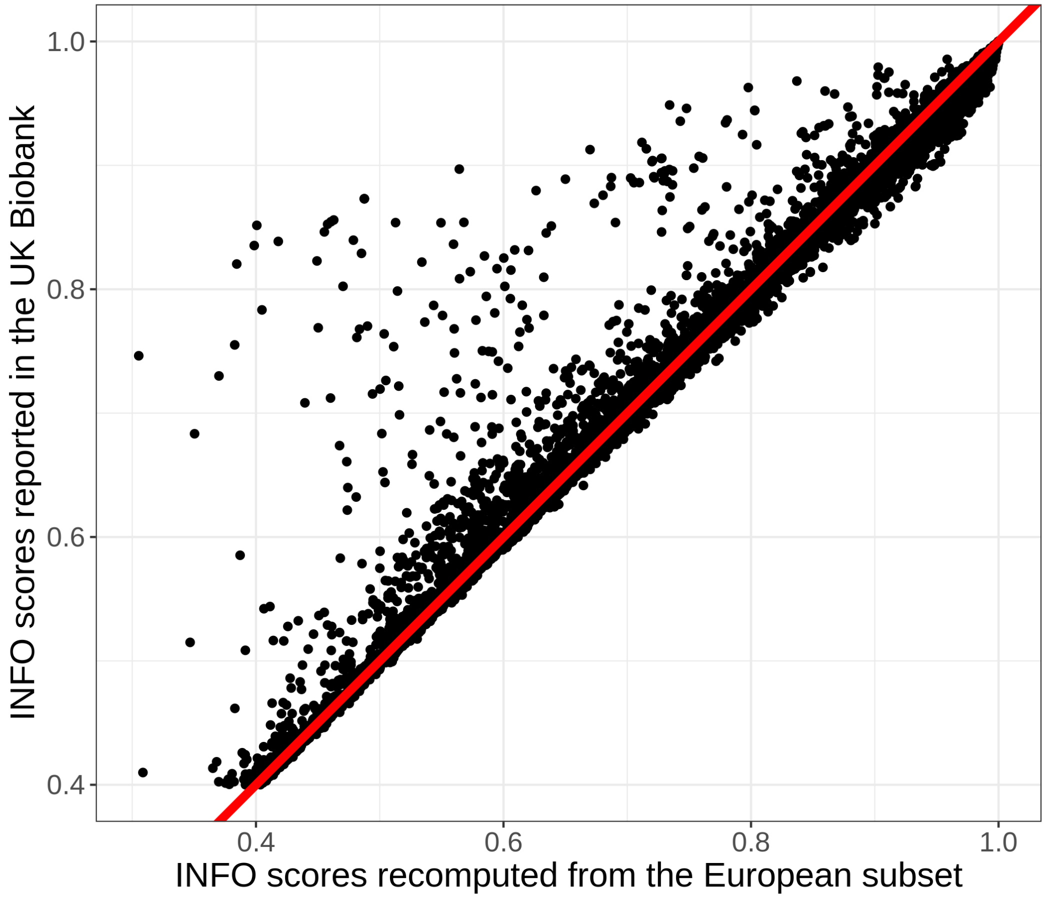


Figure S9: INFO scores reported in the UK Biobank for variants used in simulations versus INFO scores recomputed from the subset of 362,307 European individuals used in this paper.

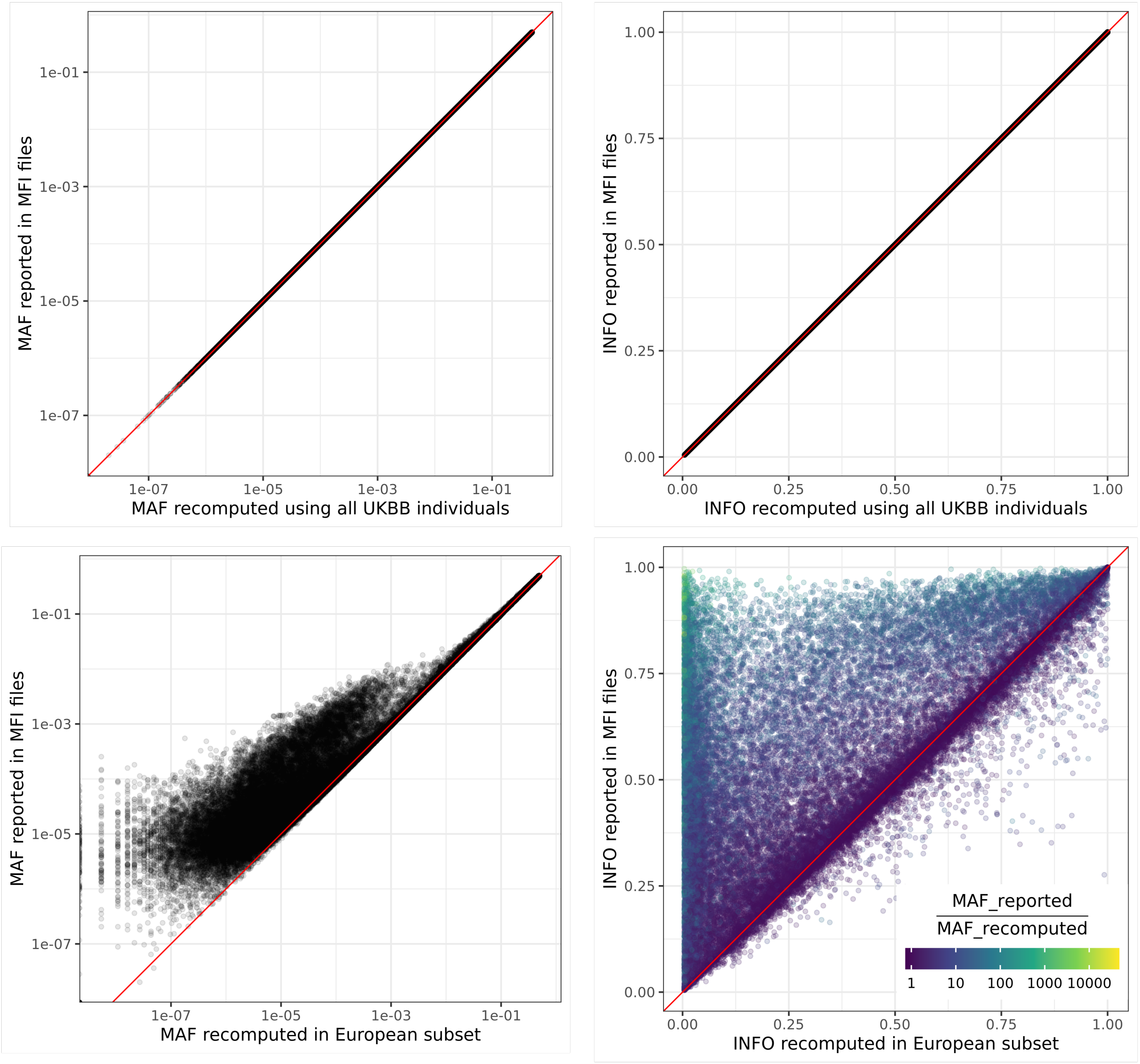


Figure S10: Minor allele frequencies (MAF) and INFO scores in the UK Biobank for 50,000 variants on chromosome 22 (chosen at random). These are either reported (in MFI files from the UK Biobank), recomputed from the whole data or from the subset of 362,307 European individuals used in this paper. MAF are represented on a log scale.

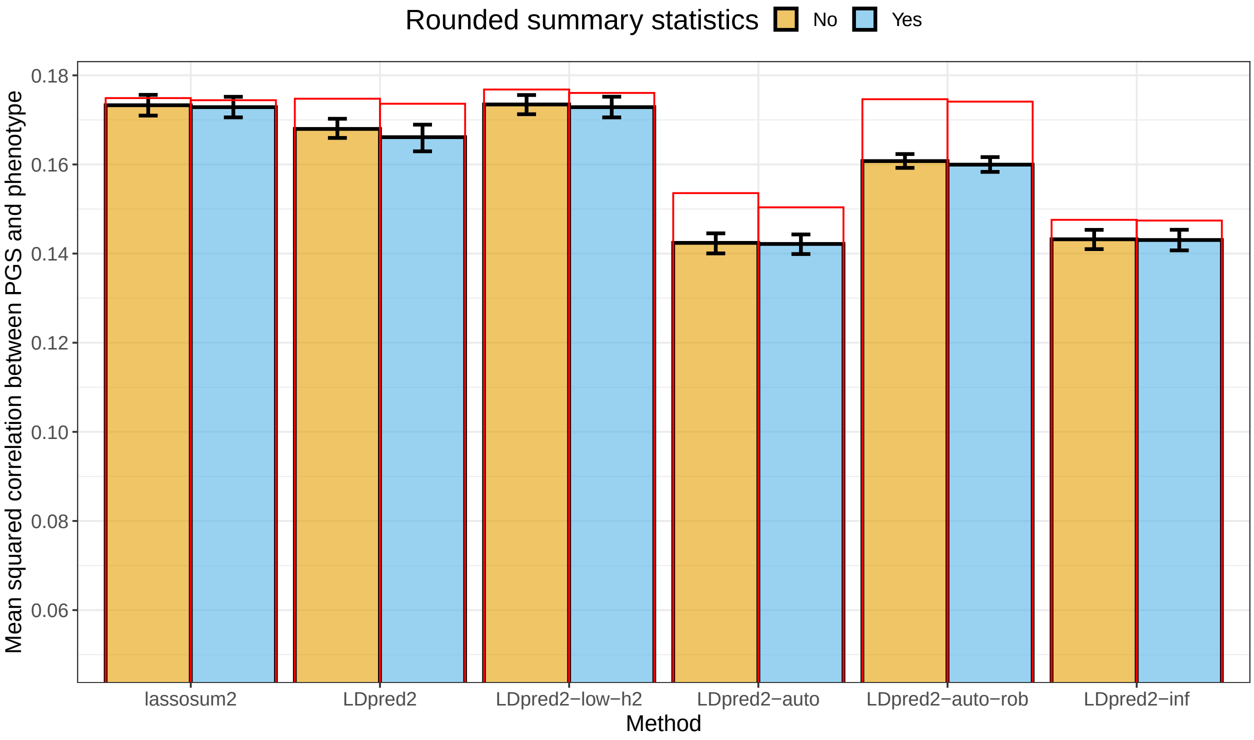


Figure S11: Results for the simulations with summary statistics (effect sizes and their standard errors) possibly rounded to two significant digits, averaged over 10 simulations for each scenario. Reported 95% confidence intervals are computed from 10,000 non-parametric bootstrap replicates of the mean. Red bars correspond to using the LD with independent blocks (Methods).

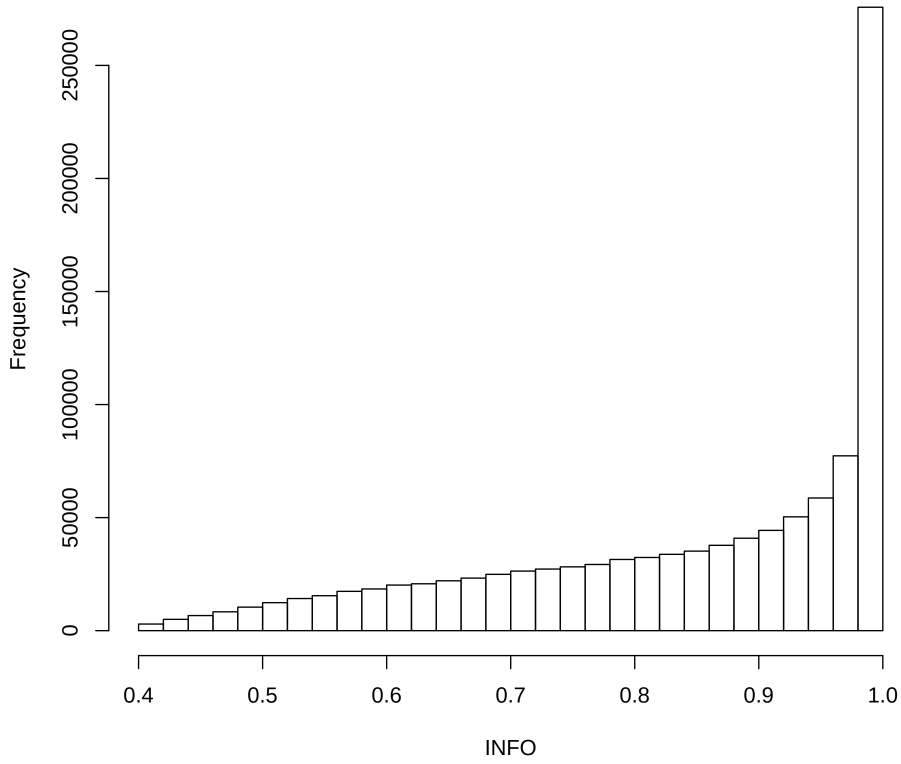


Figure S12: Histogram of the imputation INFO scores of the HapMap3 variants from the iCOGS GWAS summary statistics for breast cancer (Michailidou *et al.*, 2013).

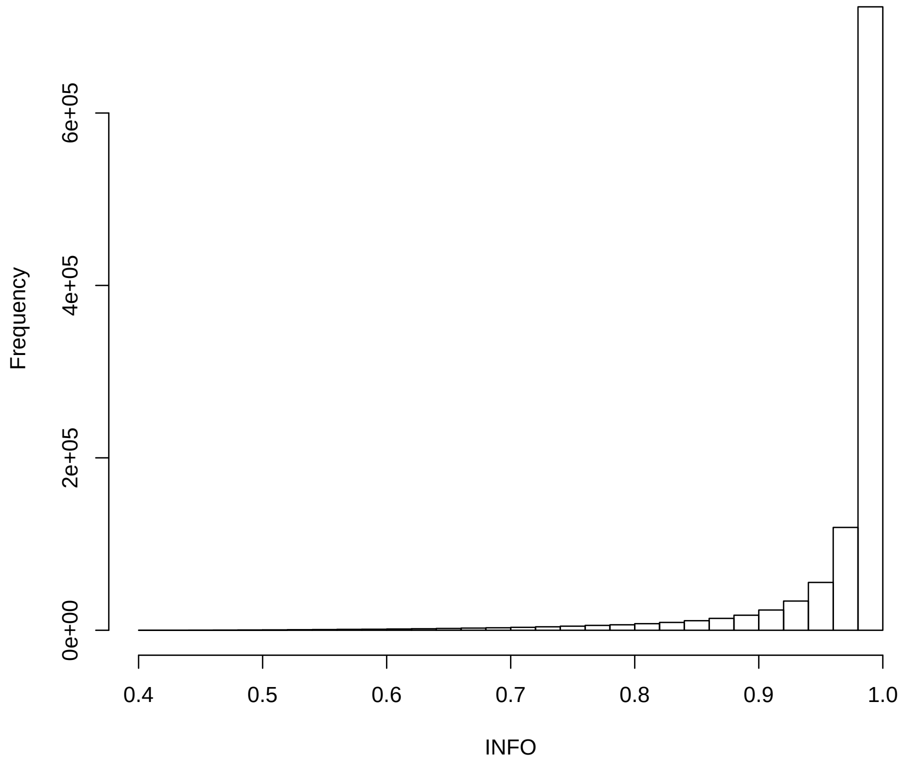


Figure S13: Histogram of the imputation INFO scores of the HapMap3 variants from the OncoArray GWAS summary statistics for breast cancer (Michailidou *et al.*, 2017).

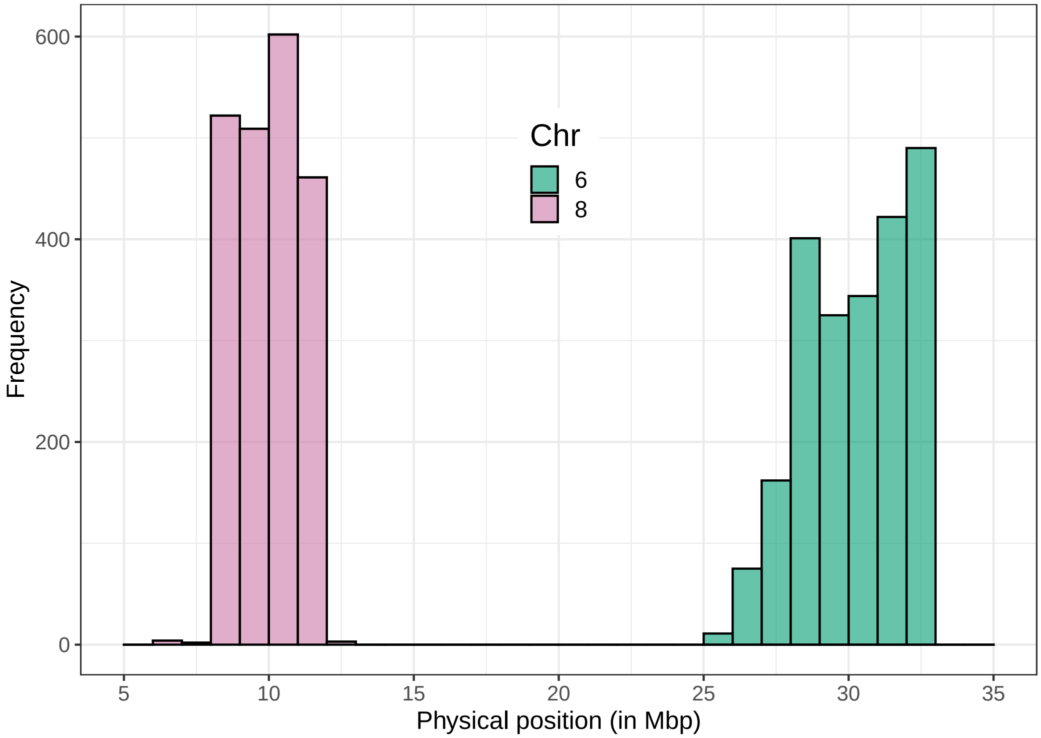


Figure S14: Histogram of the positions of outlier variants from Figure 5B in the main text.

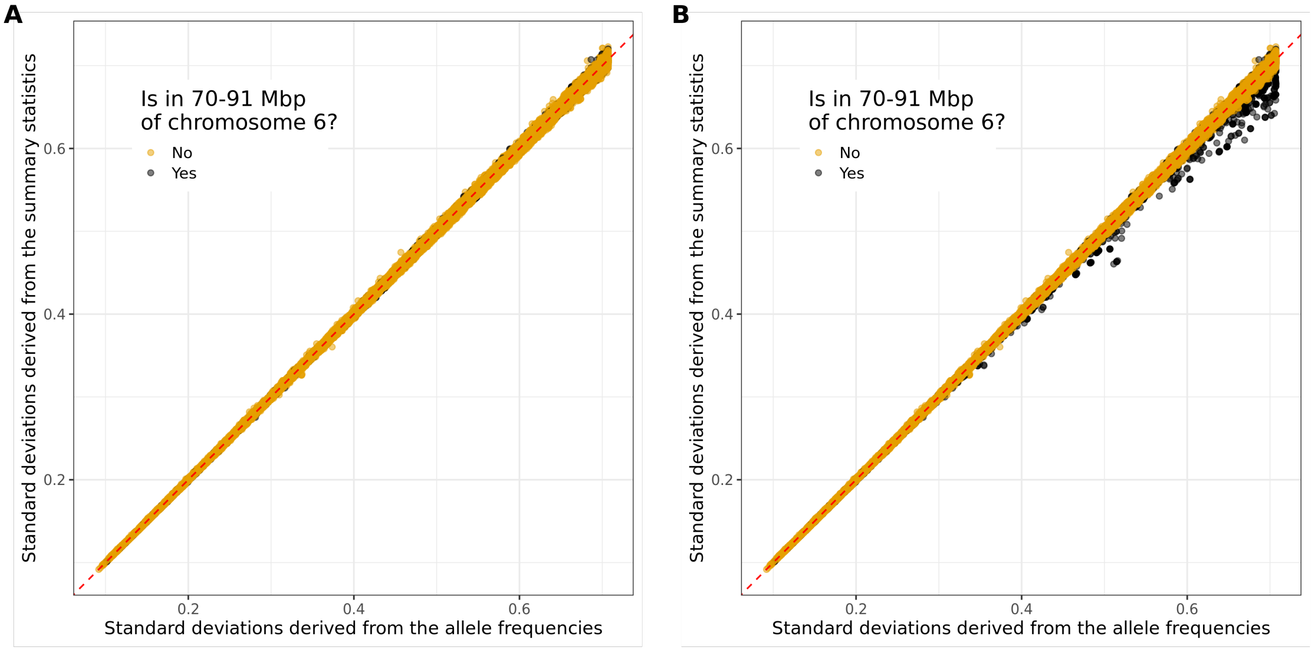


Figure S15: Standard deviations inferred from the simulated GWAS summary statistics (**A:** with no covariate; **B:** with PC19 from the UK Biobank as covariate) versus the ones inferred from the allele frequencies. Only HapMap3 variants from chromosome 6 are represented.

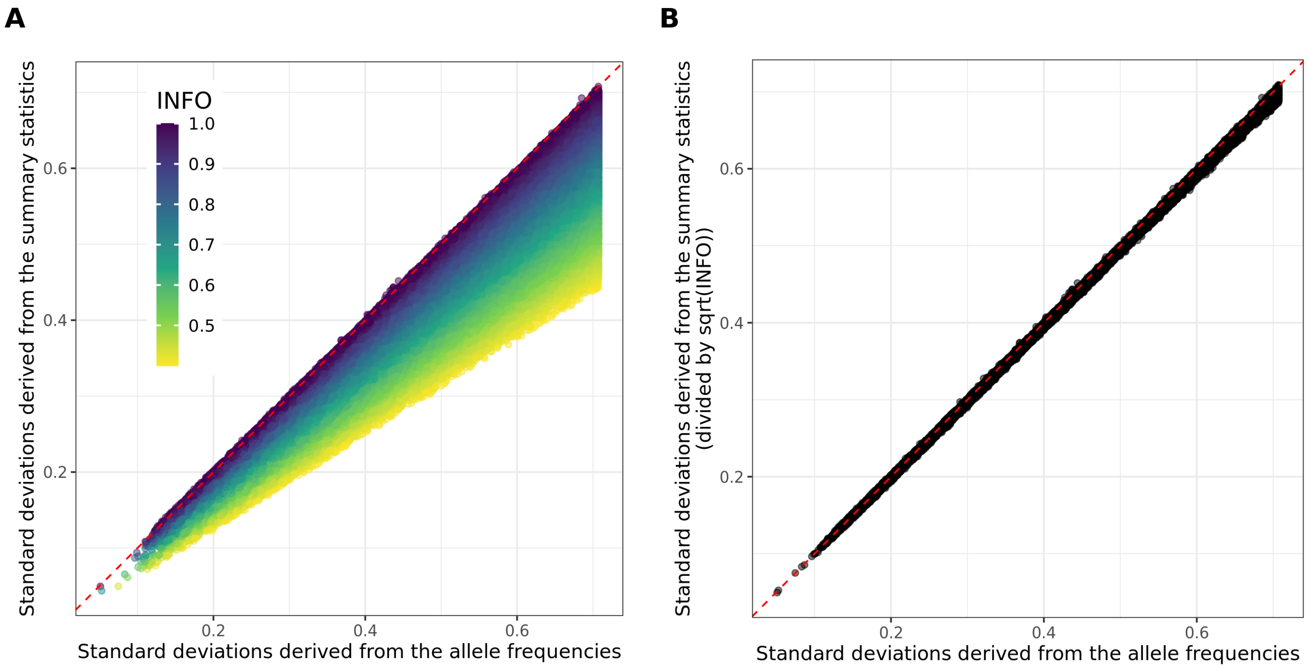


Figure S16: Standard deviations inferred from the iCOGS breast cancer GWAS summary statistics (**A:** Raw or **B:** dividing them by $\sqrt{\text{INFO}}$) versus the ones inferred from the reported GWAS allele frequencies. Only 100,000 variants are represented, at random.

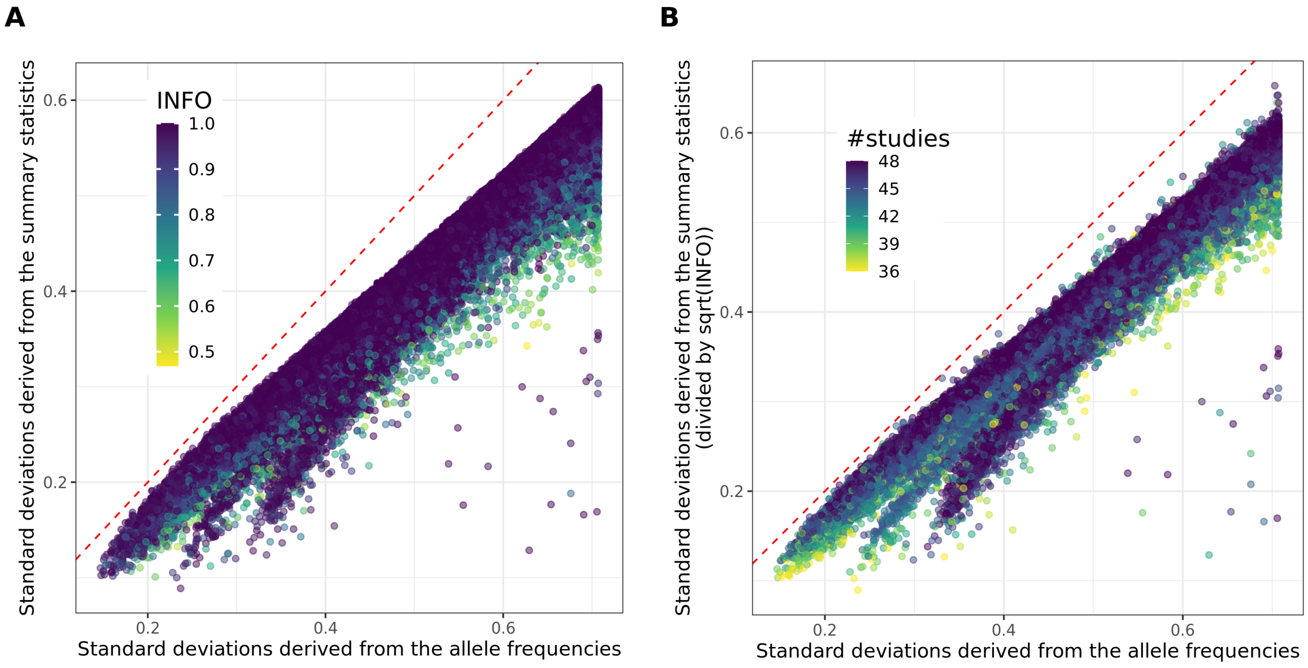


Figure S17: Standard deviations inferred from the CAD GWAS summary statistics (**A:** Raw or **B:** dividing them by $\sqrt{\text{INFO}}$) versus the ones inferred from the reported GWAS allele frequencies. Only 100,000 variants are represented, at random.

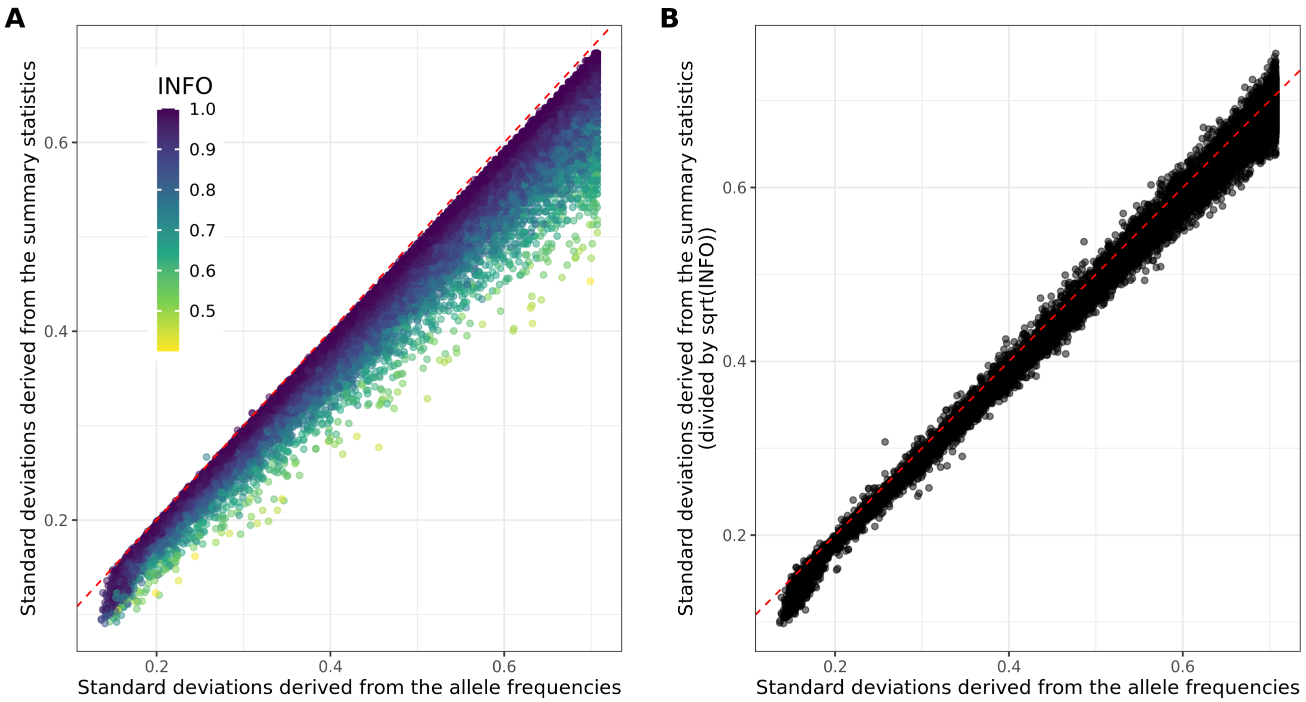


Figure S18: Standard deviations inferred from the MDD GWAS summary statistics (**A:** Raw or **B:** dividing them by $\sqrt{\text{INFO}}$) versus the ones inferred from the reported GWAS allele frequencies. Only 100,000 variants are represented, at random.

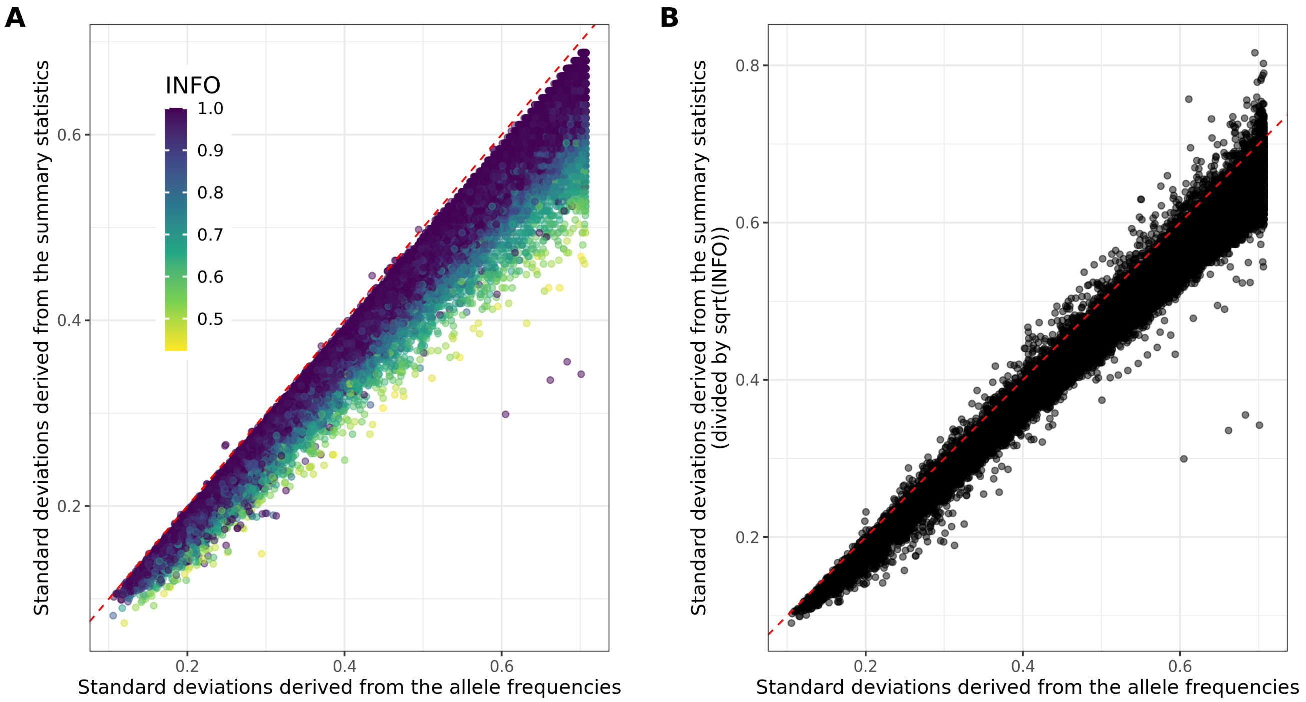


Figure S19: Standard deviations inferred from the PRCA GWAS summary statistics (**A:** Raw or **B:** dividing them by $\sqrt{\text{INFO}}$) versus the ones inferred from the reported GWAS allele frequencies. Only 100,000 variants are represented, at random.

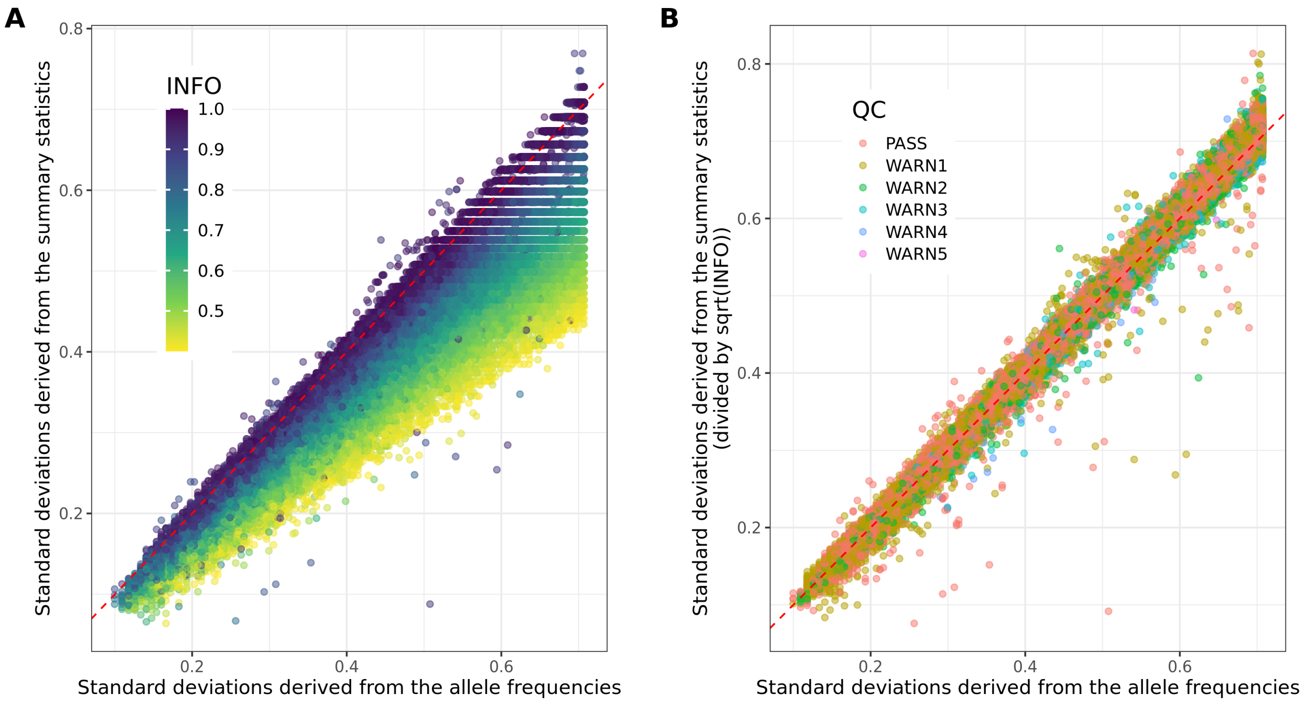


Figure S20: Standard deviations inferred from the T1D (Affymetrix) GWAS summary statistics (**A:** Raw or **B:** dividing them by $\sqrt{\text{INFO}}$) versus the ones inferred from the reported GWAS allele frequencies. Only 100,000 variants are represented, at random.

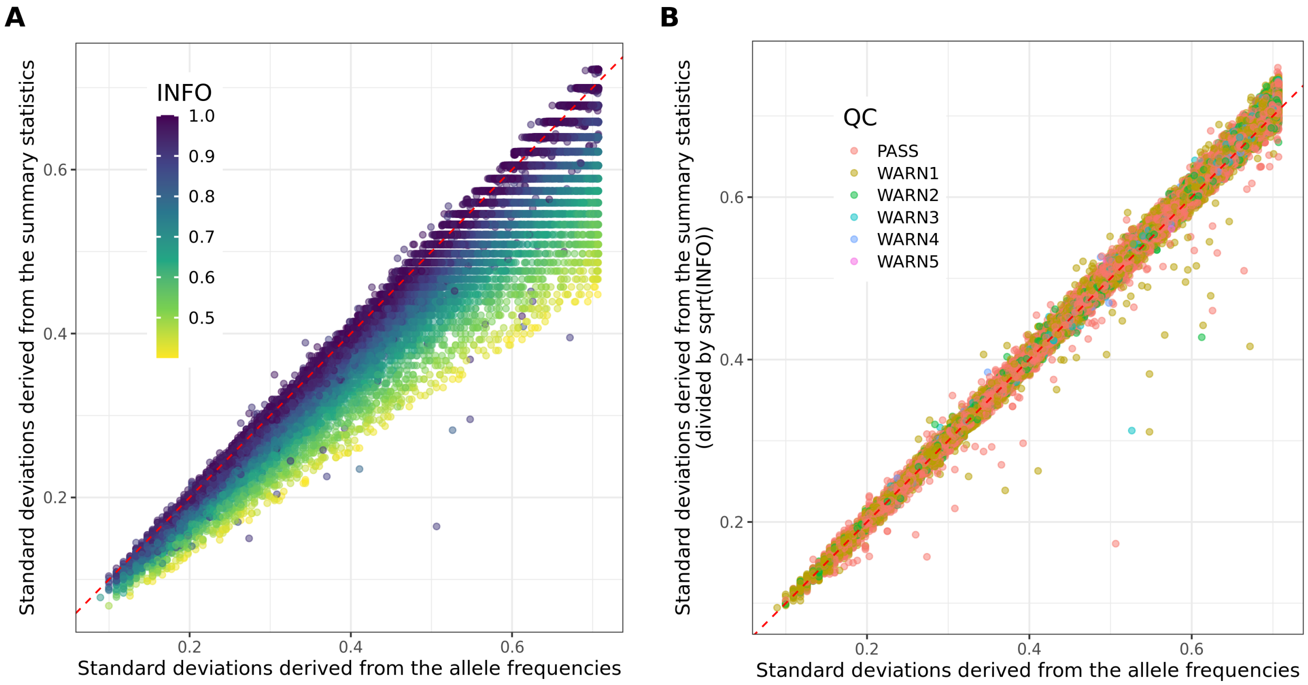


Figure S21: Standard deviations inferred from the T1D (Illumina) GWAS summary statistics (**A:** Raw or **B:** dividing them by $\sqrt{\text{INFO}}$) versus the ones inferred from the reported GWAS allele frequencies. Only 100,000 variants are represented, at random.

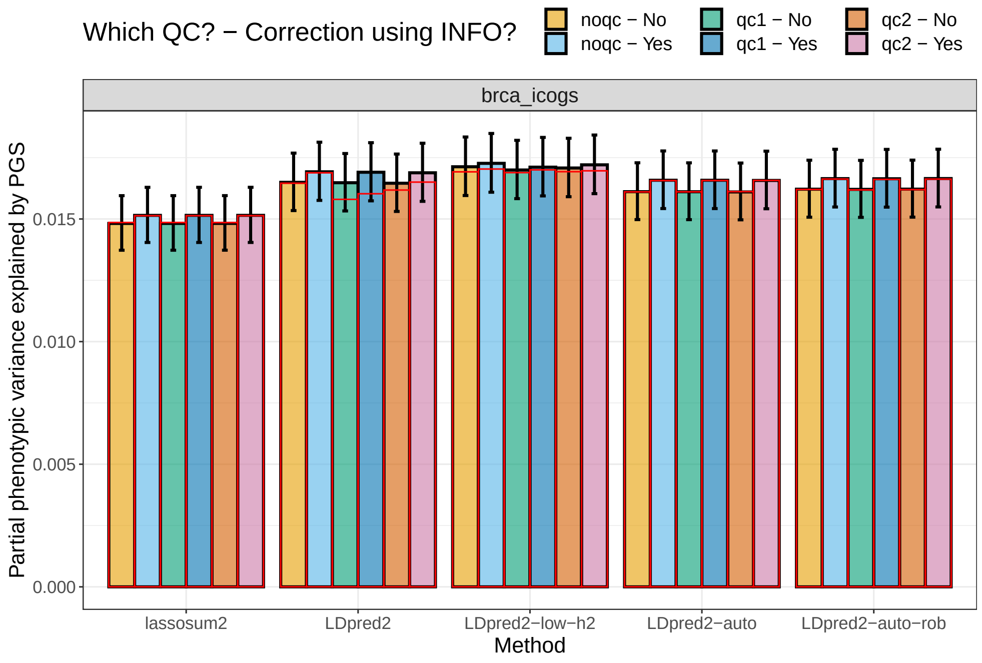


Figure S22: Variance explained of BRCA in the UK Biobank by PGS derived from external summary statistics (iCOGS). These are computed using function `pcor` of R package bigstatsr where 95% confidence intervals are obtained through Fisher's Z-transformation; these values are then squared.

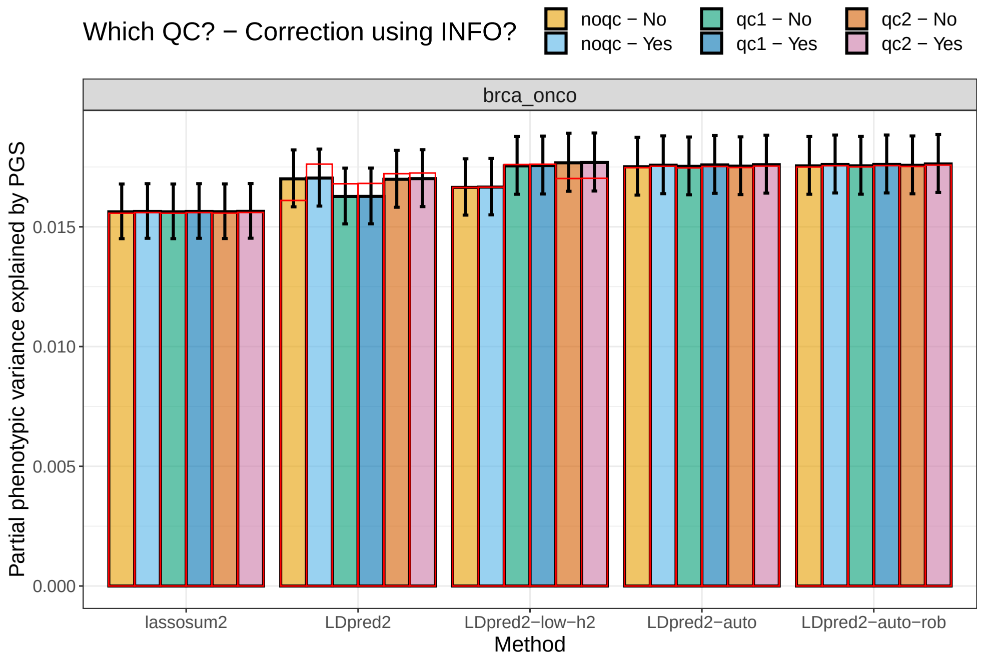


Figure S23: Variance explained of BRCA in the UK Biobank by PGS derived from external summary statistics (OncoArray). These are computed using function `pcor` of R package bigstatsr where 95% confidence intervals are obtained through Fisher's Z-transformation; these values are then squared.

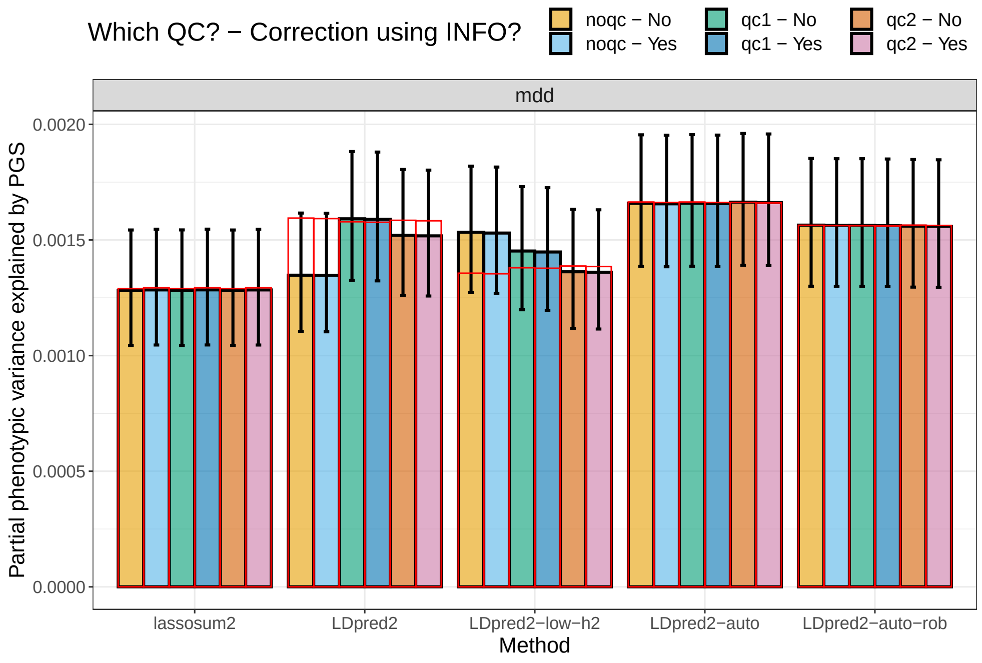


Figure S24: Variance explained of MDD in the UK Biobank by PGS derived from external summary statistics. These are computed using function `pcor` of R package bigstatsr where 95% confidence intervals are obtained through Fisher's Z-transformation; these values are then squared.

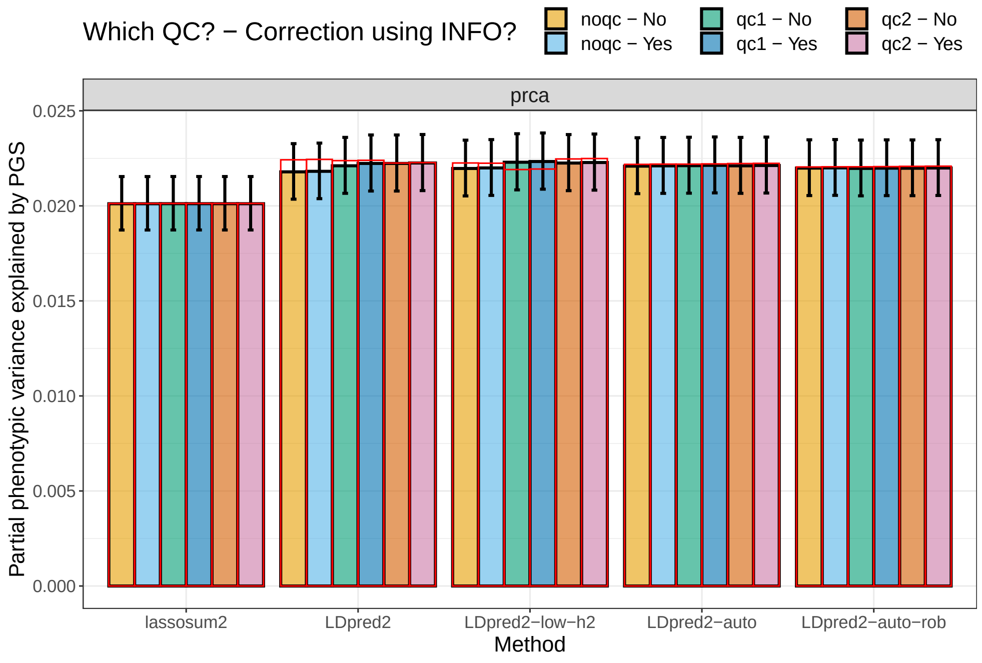


Figure S25: Variance explained of PRCA in the UK Biobank by PGS derived from external summary statistics. These are computed using function `pcor` of R package bigstatsr where 95% confidence intervals are obtained through Fisher's Z-transformation; these values are then squared.

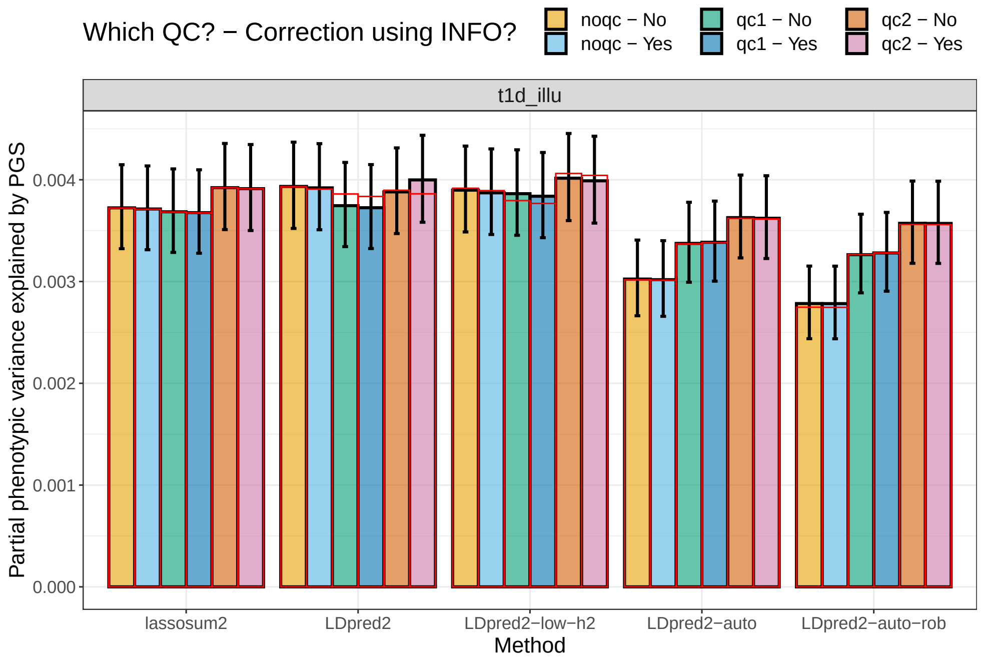


Figure S26: Variance explained of T1D in the UK Biobank by PGS derived from external summary statistics (Illumina). These are computed using function `pcor` of R package bigstatsr where 95% confidence intervals are obtained through Fisher's Z-transformation; these values are then squared.

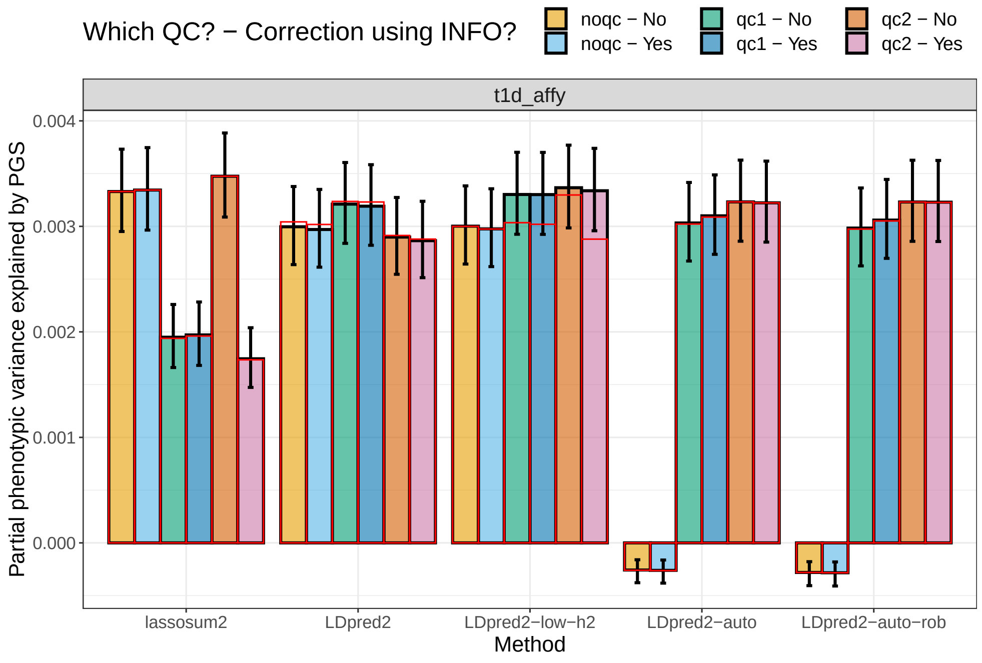


Figure S27: Variance explained of T1D in the UK Biobank by PGS derived from external summary statistics (Affymetrix). These are computed using function `pcor` of R package bigstatsr where 95% confidence intervals are obtained through Fisher's Z-transformation; these values are then squared (while keeping the sign).

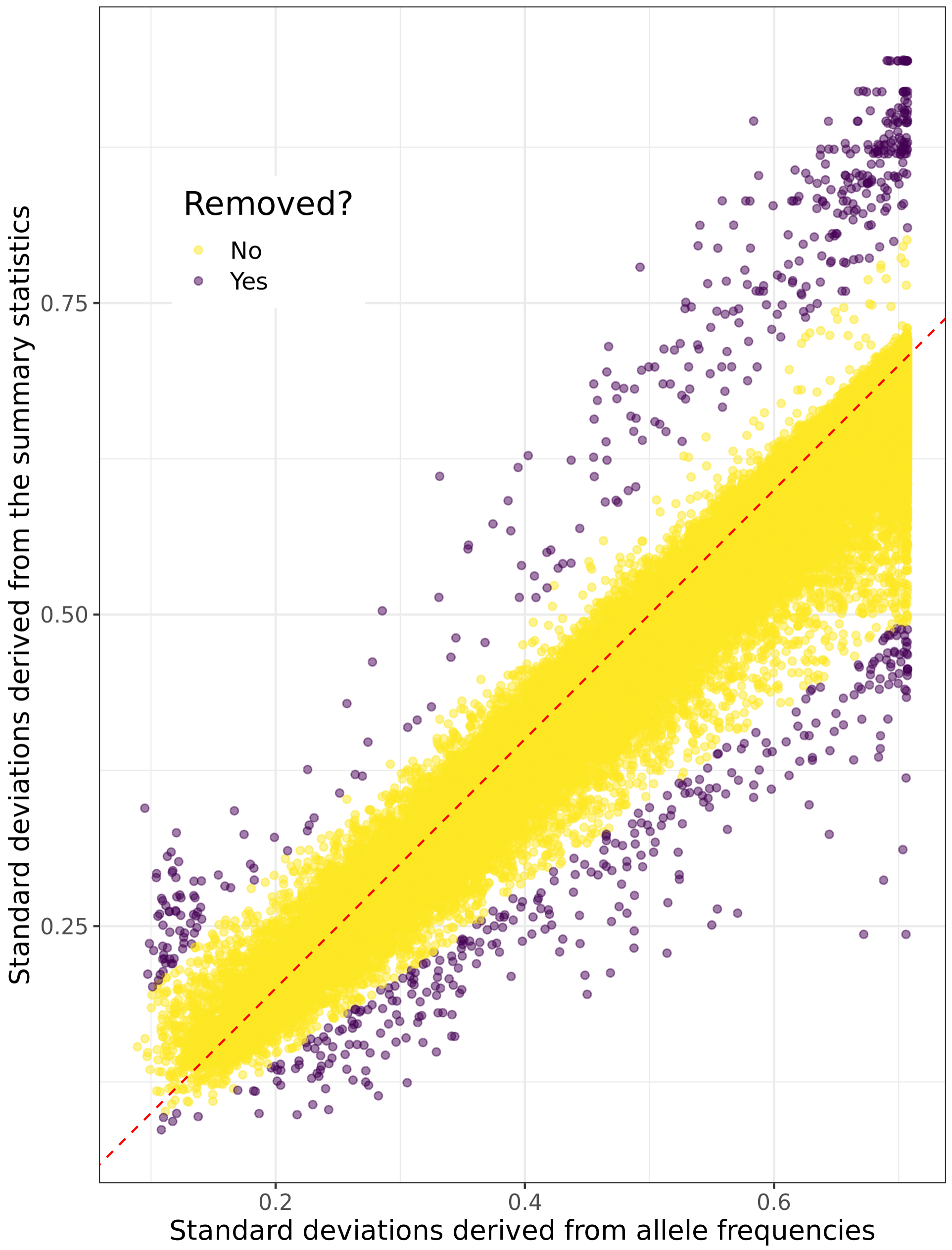


Figure S28: Standard deviations inferred from the vitamin D GWAS summary statistics versus the ones inferred from the allele frequencies of the LD reference. Only 100,000 variants are represented, at random.

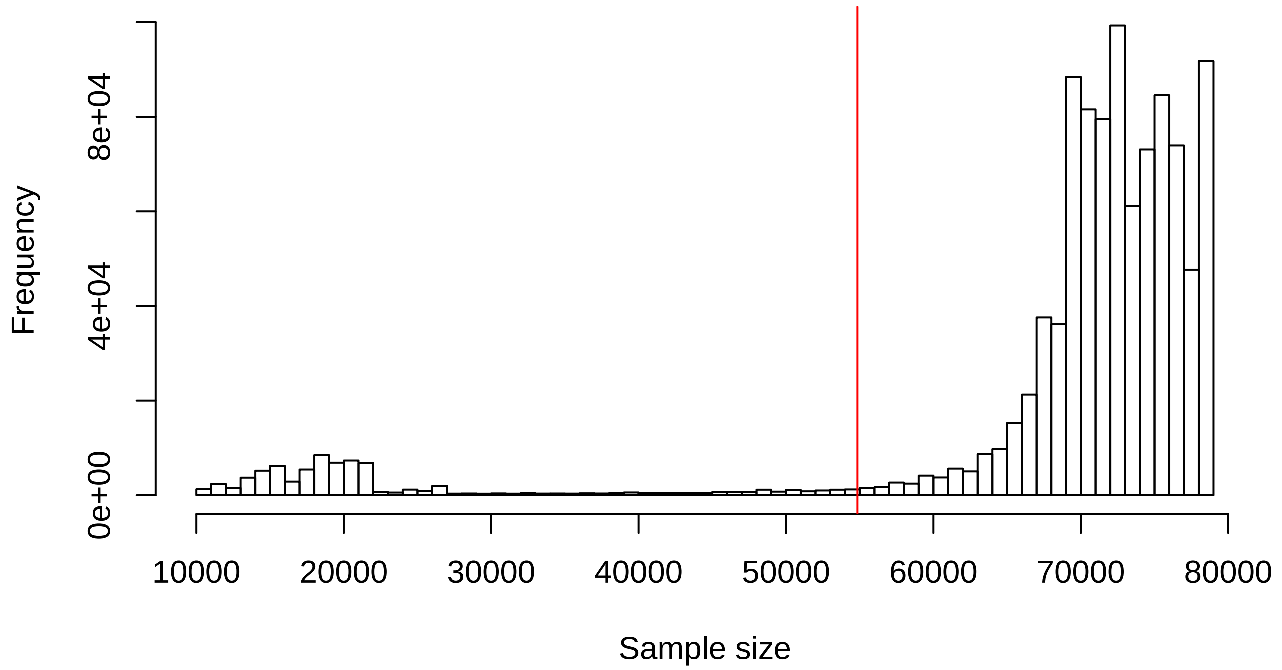


Figure S29: Histogram of the per-variant sample sizes in the vitamin D GWAS summary statistics. The vertical red line corresponds to 70% of the maximum sample size, the threshold used in “qc2”.

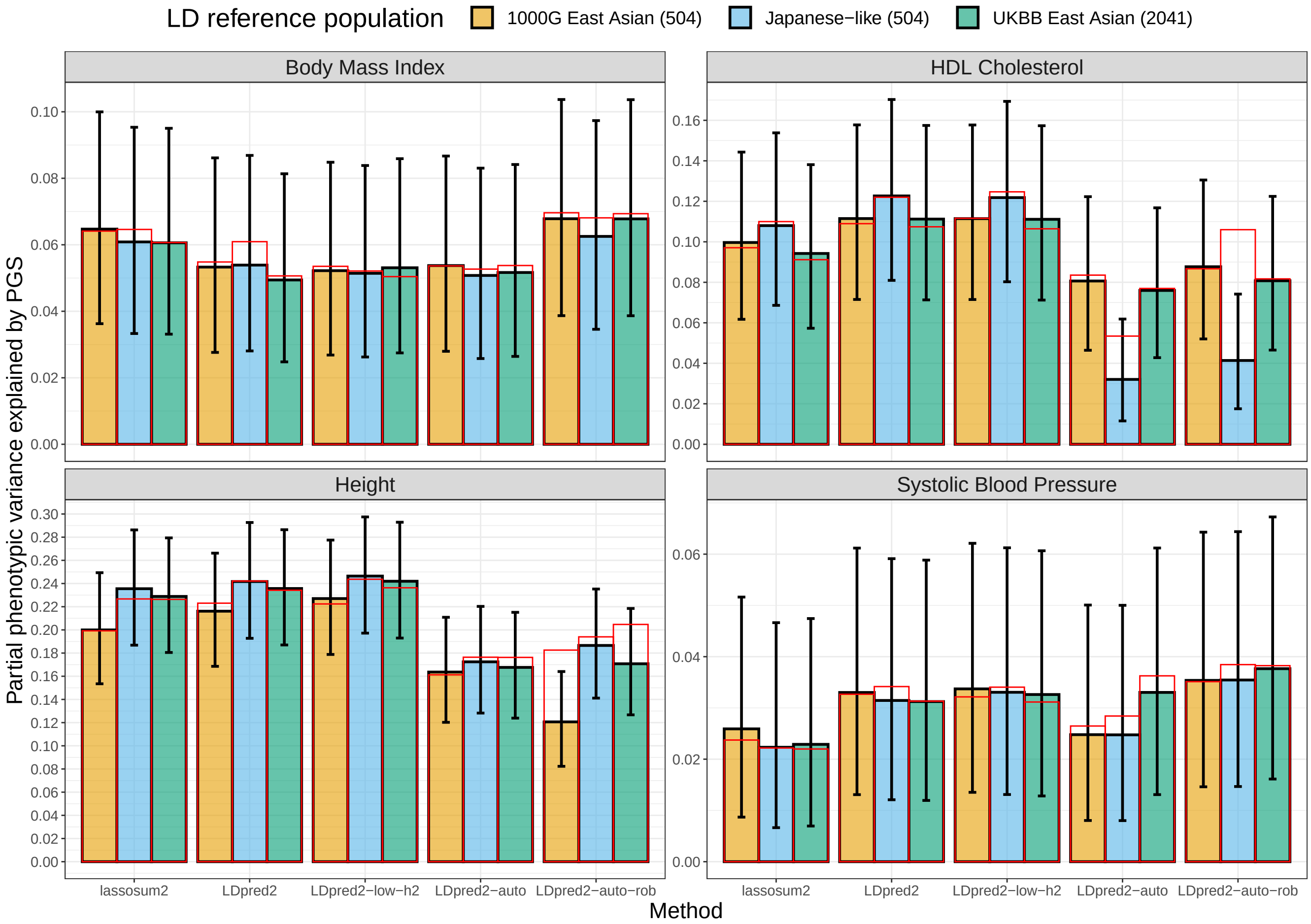


Figure S30: Results for PGS derived from four Biobank Japan GWAS summary statistics and using three different LD references. Partial correlations are computed using function `pcor` of R package bigstatsr where 95% confidence intervals are obtained through Fisher's Z-transformation, then all values are squared to report the phenotypic variance explained by PGS.

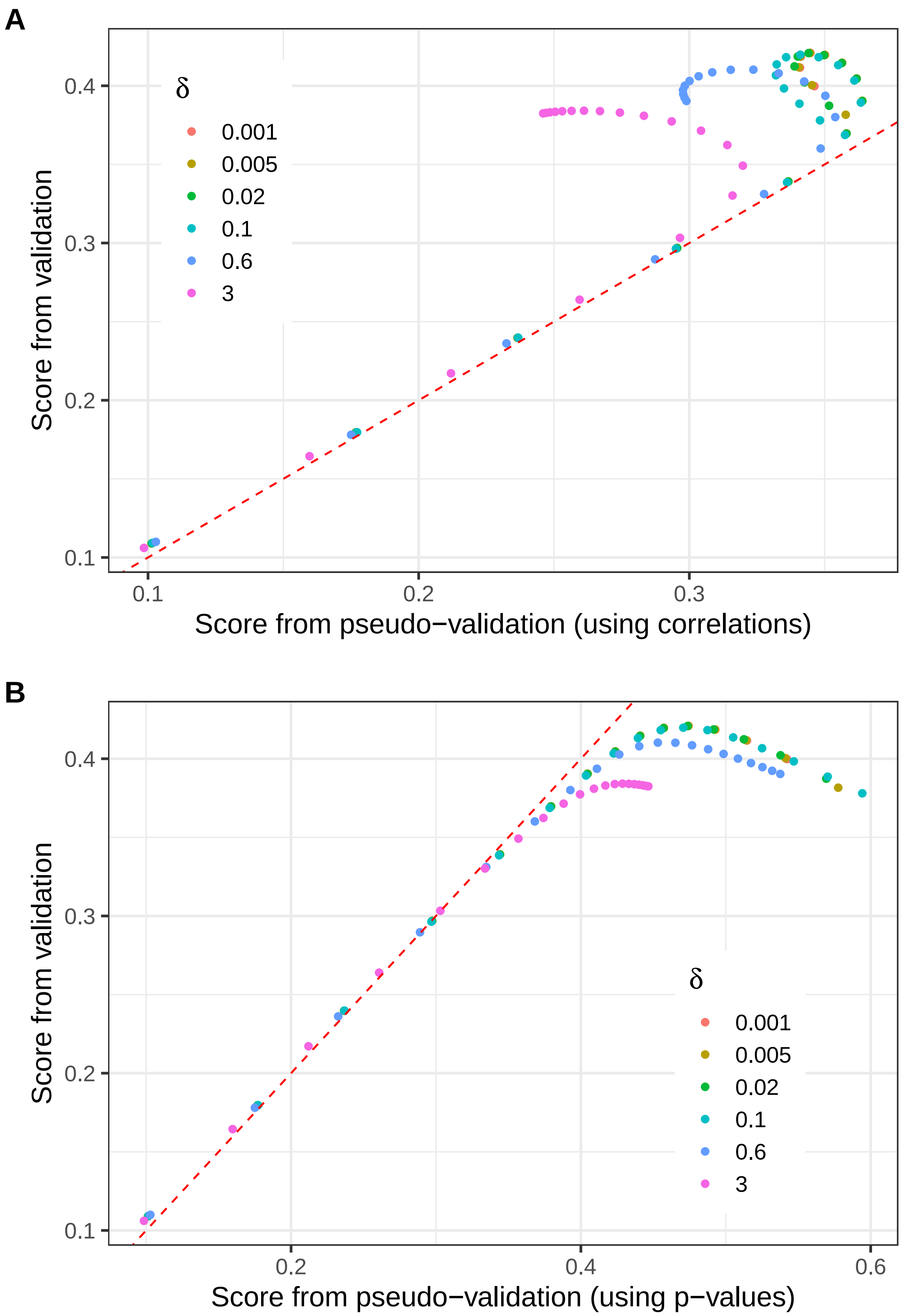


Figure S31: For one simulation, scores from validation versus scores from pseudo-validation as described in Mak *et al.* (2017) using either **A:** correlations (the default in lassosum) or **B:** p-values, when computing local false discovery rates.

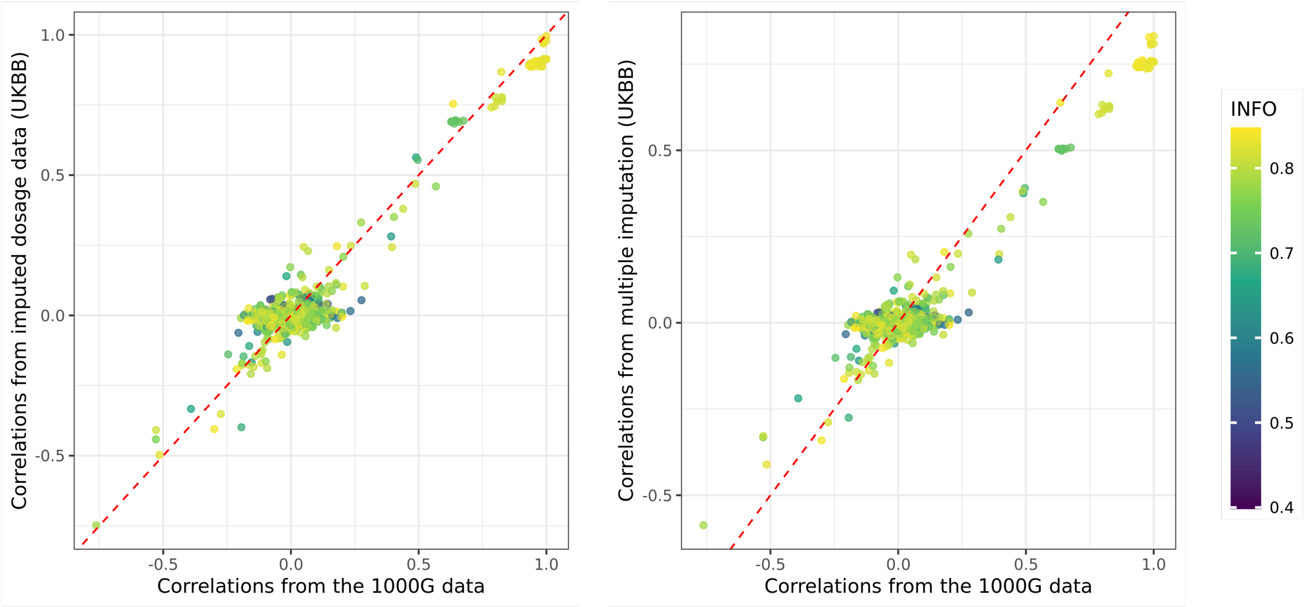


Figure S32: Comparing pairwise correlations between a subset of HapMap3 variants on chromosome 1 with an INFO score lower than 0.85, computed in three different ways: 1/ (left y-axis) from imputed dosages using 10,000 individuals from the UK Biobank (UKBB) data; 2/ (right y-axis) from multiple imputation (i.e. generating multiple complete datasets sampled according to imputation probabilities, computing correlations, and averaging results) also using UKBB; 3/ (common x-axis) from 190 individuals from the 1000 Genomes data (GBR and CEU). Each point, representing the correlation between two variants, is colored by the geometric mean of the INFO scores of these two variants.
